# Thermal Proteome Profiling reveals rapid proteomic responses to redox changes in specific cellular compartments

**DOI:** 10.1101/2024.10.17.618419

**Authors:** Aram Revazian, Alexey Nesterenko, Daria Ezeriņa, Ting Luo, Didier Vertommen, Christine S. Gibhardt, Ivan Bogeski, Joris Messens, Vsevolod V. Belousov

**Affiliations:** Molecular Physiology, Department of Cardiovascular Physiology, University Medical Center, Georg-August-University, Göttingen, Germany; Center for Precision Genome Editing and Genetic Technologies for Biomedicine, Pirogov Russian National Research Medical University, Moscow 117997, Russia; Federal Center of Brain Research and Neurotechnologies, Federal Medical Biological Agency, 117997, Moscow, Russia; VIB-VUB Center for Structural Biology, Vlaams Instituut voor Biotechnologie – Vrije Universiteit Brussel, B-1050 Brussels, Belgium; Brussels Center for Redox Biology, Vrije Universiteit Brussel, B-1050 Brussels, Belgium; Structural Biology Brussels, Vrije Universiteit Brussel, B-1050 Brussels, Belgium; de Duve Institute, Université Catholique de Louvain, 1200, Brussels, Belgium

**Keywords:** Hydrogen peroxide, HyPer-DAO, Thermal Proteome Profiling, MS-CETSA, Redox-signaling, Reactive Oxygen Species

## Abstract

Hydrogen peroxide (H_2_O_2_) functions as a secondary messenger in cellular redox signaling, acting mainly via oxidation of protein thiols. Its spatially and temporally regulated activity within cells is essential for maintaining proper redox balance, and disruptions in these patterns can lead to oxidative stress and various related pathologies. Redox proteomics, which examines the impact of H_2_O_2_ at the proteome level, typically focuses only on thiol oxidation, overlooking broader proteomic alterations and the significance of subcellular localization in these redox processes. In this study, we address these open questions by combining chemogenetics with Thermal Proteome Profiling (TPP) to map global proteome response to compartmentalized H_2_O_2_ production. We identified hundreds of proteins with altered thermostability and/or abundance upon localized H_2_O_2_ generation in the cytosol, nucleus, and the ER lumen, highlighting their roles in cellular responses to localized H_2_O_2_. We identified proteins such as MAP2K1, PARK7, TRAP1, and UBA2 to be highly sensitive to localized H_2_O_2_ production. Furthermore, we validated their altered thermostability and found that these changes are controlled via dysregulated protein-protein interactions. This study provides a valuable resource for researchers exploring redox-mediated signal transduction and offers novel insights that could be harnessed in treating oxidative stress-induced pathologies.

## Introduction

Reactive oxygen species (ROS) is a common term widely used to describe various partially reduced oxygen derivatives. Initially viewed merely as unavoidable by-products of aerobic life to be eliminated at all costs ^1^, ROS can damage a wide area of biological molecules such as DNA, lipids, proteins^2^, contributing to various disorders like ageing, cancer, neurodegenerative and cardiovascular diseases^3^. However, over the past few decades, this perception has evolved with the recognition of at least some ROS as signaling molecules for inter–^4^ and intracellular communication^5^. ROS can manifest as radicals, non-radicals, or electrically charged or uncharged entities, each possessing distinct thermodynamic reduction-oxidation (redox) potentials and kinetic properties^1^. As a result, biologically relevant ROS exhibit varying abilities to diffuse, penetrate lipid membranes, and interact with biological molecules. Among ROS, hydrogen peroxide (H_2_O_2_) stands out as the most extensively studied, boasting properties well-suited for signaling functions.

Although H_2_O_2_ is a potent oxidant, its reactivity with most biomolecules including low molecular weight antioxidants and most protein thiols^6^, is limited. Except for a few exceptions, thiols react slowly with H_2_O_2_, typically only in the form of a thiolate anion (deprotonated SH-group). H_2_O_2_-mediated redox signaling primarily involves the formation of oxidative post-translational modifications of cysteines in proteins, altering protein activity, conformation, and function. Due to its relatively low reactivity, H_2_O_2_ can potentially diffuse over considerable distances in aqueous environments^1^. However, the cell’s abundant expression of peroxidases, such as peroxiredoxins and glutathione peroxidases, acts as sinks for H_2_O_2_, thereby dampening (or terminating) H_2_O_2_-mediated redox signaling^7^. Furthermore, the cell can regulate H_2_O_2_ transport across biological membranes via lipid composition and membrane channels like aquaporins^8^, providing another mechanism to control the propagation of redox signals in a spatiotemporal manner.

The compartmentalization of eukaryotic cells into organelles allows for the spatial separation of distinct processes and the establishment of optimal conditions for each. Consequently, different cellular compartments contain a heterogeneous mix of pro– and antioxidant systems, as well as redox-sensitive proteins^9–12^. The global cellular response to an increase in H_2_O_2_ within a specific organelle depends on the abundance of antioxidant systems in each compartment and the ability of H_2_O_2_ to cross the membrane of the organelle producing it. Most studies that add H_2_O_2_ externally (e.g., as a bolus) fail to replicate physiologically relevant redox changes, which often originate from specific organelles^13^.

To address this issue, several strategies have been developed to modulate endogenous H_2_O_2_ levels. Depending on the target organelle, ROS levels can be adjusted by regulating the copy number of a specific organelle, modulating the activity of an H_2_O_2_-coupled metabolic process within the organelle of interest, or adjusting the activity of endogenous pro-or antioxidant enzymes^14^. However, these approaches do not allow precise control over H_2_O_2_ production because of very complex regulation of the activity of these sources.

Alternatively, ROS can be manipulated in cells at the level of cellular compartments and sub-compartments using optogenetic and chemogenetic approaches. In the first case, genetically encoded photosensitizers can produce ROS upon light irradiation^15,16^. In the second case, organelle-specific H_2_O_2_ generation can be achieved using enzymes metabolizing specific substrates^17^. In our study, we utilized D-amino acid oxidase (DAO), an enzyme that produces H_2_O_2_ in response to D-amino acids, expressed in the cytosol, nucleus, and ER lumen of cells^18^. This approach allowed us to generate H_2_O_2_ endogenously and study its compartment-specific effects.

Although it is already challenging to reconstitute redox alterations akin to those observed in cells under physiological or pathological conditions, yet, an even greater challenge lies in identifying the proteins involved in cellular response to these alterations. Given the complexity and multifaceted nature of cellular responses to redox perturbations, proteomic methods, primarily based on mass spectrometry (MS) for detection, have become indispensable tools in studying cellular redox signaling. While various redox proteomic methods have been developed^19–24^, most focus on techniques on capturing the state of protein cysteine thiols^25^, thereby limiting the identification of downstream participants of redox signaling pathways that are not directly modified by the redox agent (via non-redox mechanisms).

Thermal Proteome Profiling (TPP), also known as MS-CETSA, offers a quantitative proteomics approach, which is based on the tendency of proteins to denature, aggregate and precipitate upon heating, enabling the study of rapid cellular responses to various stimuli^26,27^. TPP evaluates the “thermostability” of proteins, influenced by interactions with other proteins, ligand binding, post-translational modifications (PTMs), and other factors^28^. In contrast to conventional proteomics which detects mostly alterations of the protein levels caused by protein synthesis and degradation, TPP can measure acute response of the proteome to certain stimuli. Unlike methods solely targeting direct redox targets, TPP can assess all identified proteins for altered thermostability, providing a powerful alternative or complementary approach to redox proteomics for investigating cellular response to redox alterations.

In our study, we employed multiple methodologies to address the limitations of many redox protein studies. To induce organelle-specific H_2_O_2_ generation, we utilized DAO expressed in the cytosol, nucleus, and ER lumen of living cells. To identify proteins involved in the acute cellular response to this organelle-specific H_2_O_2_ generation independently of thiol states, we applied the TPP method. This led to the identification of several hundred proteins with altered thermostability and/or abundance in at least one selected organelle. We then selected four proteins — MAP2K1, PARK7, TRAP1, and UBA2 — as bait proteins in co-immunoprecipitation experiments to determine whether changes in their thermostability, as identified by TPP, are influenced by differences in their protein-protein interaction networks. While some proteins may have undergone thermostability shifts due to H_2_O_2_-induced changes in their interactome, other factors appear to be at play for certain proteins.

## Results

### Creating cell lines for compartment-specific H₂O₂ generation

The significance of compartment-specific redox signaling in cells offers opportunities for developing biological models, enabling local manipulation and real-time monitoring of redox processes. Previous studies have successfully utilized chemogenetic H_2_O_2_ generation by yeast DAO in eukaryotic cells, in combination with the HyPer family probes, to visualise H_2_O_2_ production^29,30^. In our study, we constructed three vectors encoding the HyPer-DAO fusion for three cellular compartments: the cytosol (pLVX-Puro-HyPer-DAO-NES), the nucleus (pLVX-Puro-HyPer-DAO-3NLS), and the endoplasmic reticulum (ER) lumen (pLVX-Puro-TagBFP-DAO-KDEL). Each plasmid contains a CMV promoter, a fluorescent protein, DAO, and a compartment-targeting signal. For the ER-targeted plasmid, we substituted HyPer with TagBFP, as HyPer becomes fully oxidised in this compartment^11,31^. These plasmids were then used to produce lentiviruses and establish stable HEK293 cell lines (cyto-DAO, nuc-DAO and ER-DAO) expressing HyPer-DAO or TagBFP-DAO proteins in the respective cellular compartments. Cell lines expressing HyPer-DAO were subjected to monoclonal selection (Figure 1A).

**Figure 1.**
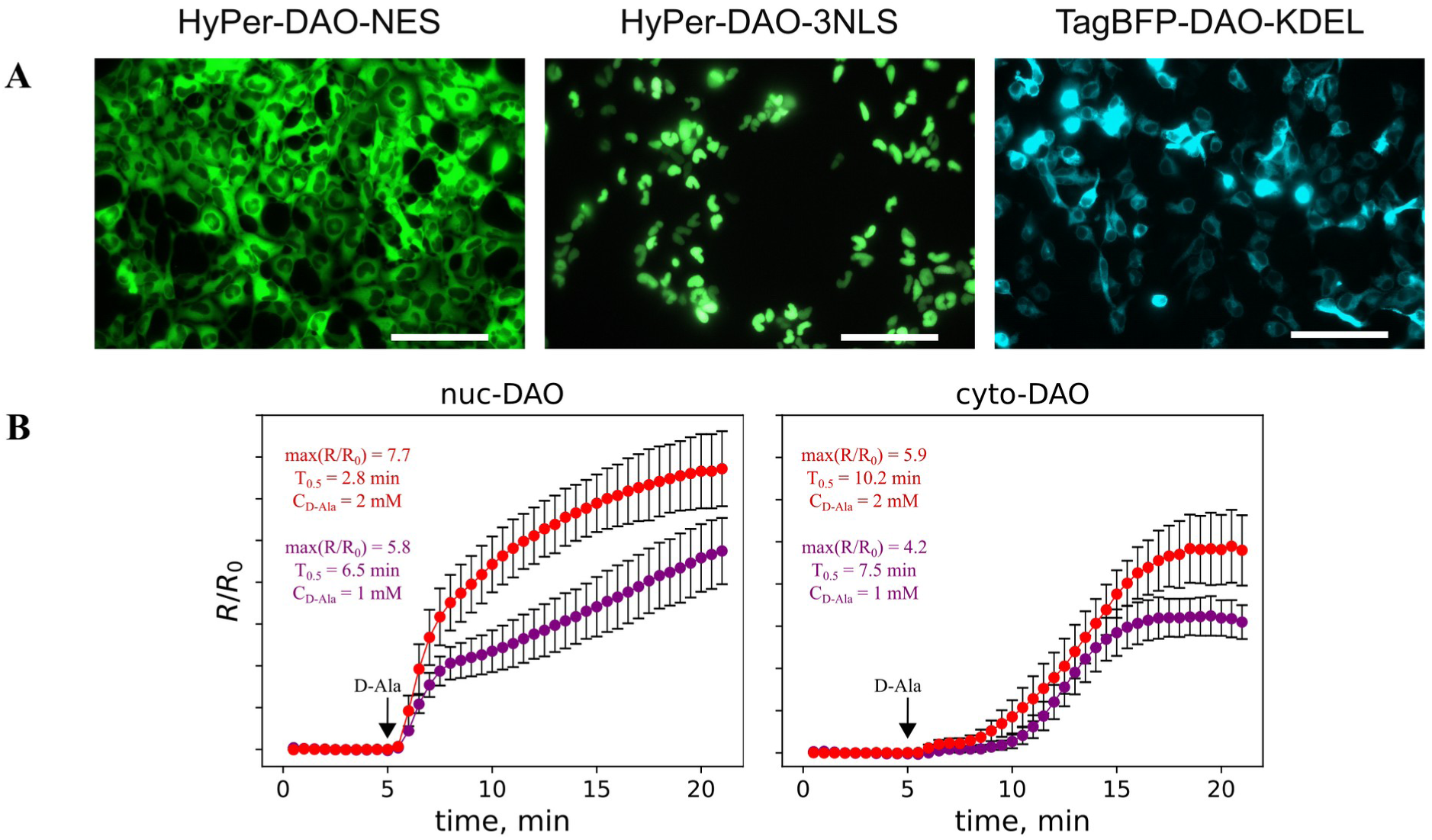
DAO-driven compartment-specific H_2_O_2_ generation. (A) Stable HEK293 cell lines, expressing HyPer-DAO or TagBFP-DAO chimeric proteins in the cell cytosol (cyto-DAO), nucleus (nuc-DAO) or ER lumen (ER-DAO). Scale bar 100 μm. (B) HyPer oxidation kinetics in response to addition of D-Ala to cyto-DAO and nuc-DAO cell lines. R/R_0_ – normalized HyPer ratio signal. Max(R/R_0_) – maximal value of the HyPer ratio signal it reaches upon addition of D-Ala during the experiment. T_0.5_ – the time required for HyPer to reach the half of the maximal ratio. Each curve represents the mean response of at least 10 cells from the same dish, bars represent standard deviation (SD).

We then optimized the D-Alanine (D-Ala) concentration required for DAO stimulation in these cell lines. A concentration of 1 mM D-Ala induced near-maximal oxidation of HyPer in both cyto-DAO and nuc-DAO cell lines (Figure 1B). Since HyPer is fused to DAO, H_2_O_2_ levels in the immediate vicinity of HyPer are assumed to be the highest, compared to more distant locations within the same organelle. Therefore, to ensure robust elevation of H_2_O_2_ within the organelle of interest, we decided to use 2 mM D-Ala for DAO stimulation in both cytosolic and nuclear cell lines. As TagBFP cannot detect H_2_O_2_ in the ER lumen upon addition of D-Ala, and assuming that the ER membrane may act as a barrier for D-Ala transport, we decided to use 8 mM D-Ala in the ER-DAO cell line.

### Thermal proteome profiling of cells upon compartmentalised H_2_O_2_ generation

To identify proteins involved in the cellular response to H_2_O_2_, independent of redox-sensitive cysteines in their amino-acid sequences, we utilized Thermal Proteome Profiling (TPP). TPP is based on the principle that changes in the state of protein impact its thermal denaturation profile, referred to as the “melting curve”. By comparing the melting curves of proteins under experimental and control conditions, TPP detects changes in protein thermostability.

Details of TPP procedure are described in the method section (Figure S1). The raw mass spectrometric (MS) data were processed, resulting in the identification of 5,252 proteins, each representing at least two peptides. This substantial number, alongside the high degree of overlap with other recent TPP studies ^32–34^, suggests comprehensive proteome coverage (Figure S2A, Supplementary Note S1).

To compare the responses of different cell lines to H_2_O_2_, we multiplexed cell lines and conditions in a single MS run using the 2D-TPP approach ^35^. This multiplexing strategy ensures consistent identification (or exclusion) of proteins across all conditions. The 2D-TPP method allowed us to calculate ratios of experimental-to-control protein quantities at specific temperatures, based on two TPP metrics: abundance and stability scores^34^. The abundance score reflects changes in protein levels in the soluble fraction at the lowest treatment temperatures, while the stability score measures resistance to heat-induced denaturation (Figure S2B). Proteins were classified as “hits” or “no hits” based on the fold changes in stability and abundance scores, along with statistical significance (adjusted p-value). The complete list of proteins is supplemented (Table S1). As expected, very few hits were identified in the wild-type cell line, which cannot generate H_2_O_2_ in response to D-Ala. In contrast, a higher number of hits were found in the ER-DAO cell line upon H_2_O_2_ generation, with even more identified in the nuc-DAO cell line, and the most in the cyto-DAO cell line (Figure S2C-D).

### System analysis of thermoproteome changes

To identify patterns in protein distribution based on their response to H_2_O_2_, we performed a cluster analysis (Figure 2). Proteins were grouped based on their abundance and stability scores for each of four cell lines. These clusters were then categorized into three groups based on the abundance and stability scores of centroid proteins within each cluster. The first group included six clusters, where the centroid proteins displayed similar abundance and stability scores in response to H_2_O_2_ generated in the cytosol and the nucleus, with significantly higher scores compared to H_2_O_2_ generation in the ER lumen (Figure 2). The second group consisted of two clusters and contained centroid proteins with more pronounced changes in thermostability (but not abundance) in response to H_2_O_2_ generated in the cytosol than in either the nucleus or the ER lumen. The third group included only one cluster with 12 proteins (Figure 2). The centroid protein of this cluster exhibited high absolute abundance and stability scores when H_2_O_2_was produced in the ER-DAO cell line, with relatively less pronounced changes in the nuc-DAO and cyto-DAO cell lines.

**Figure 2.**
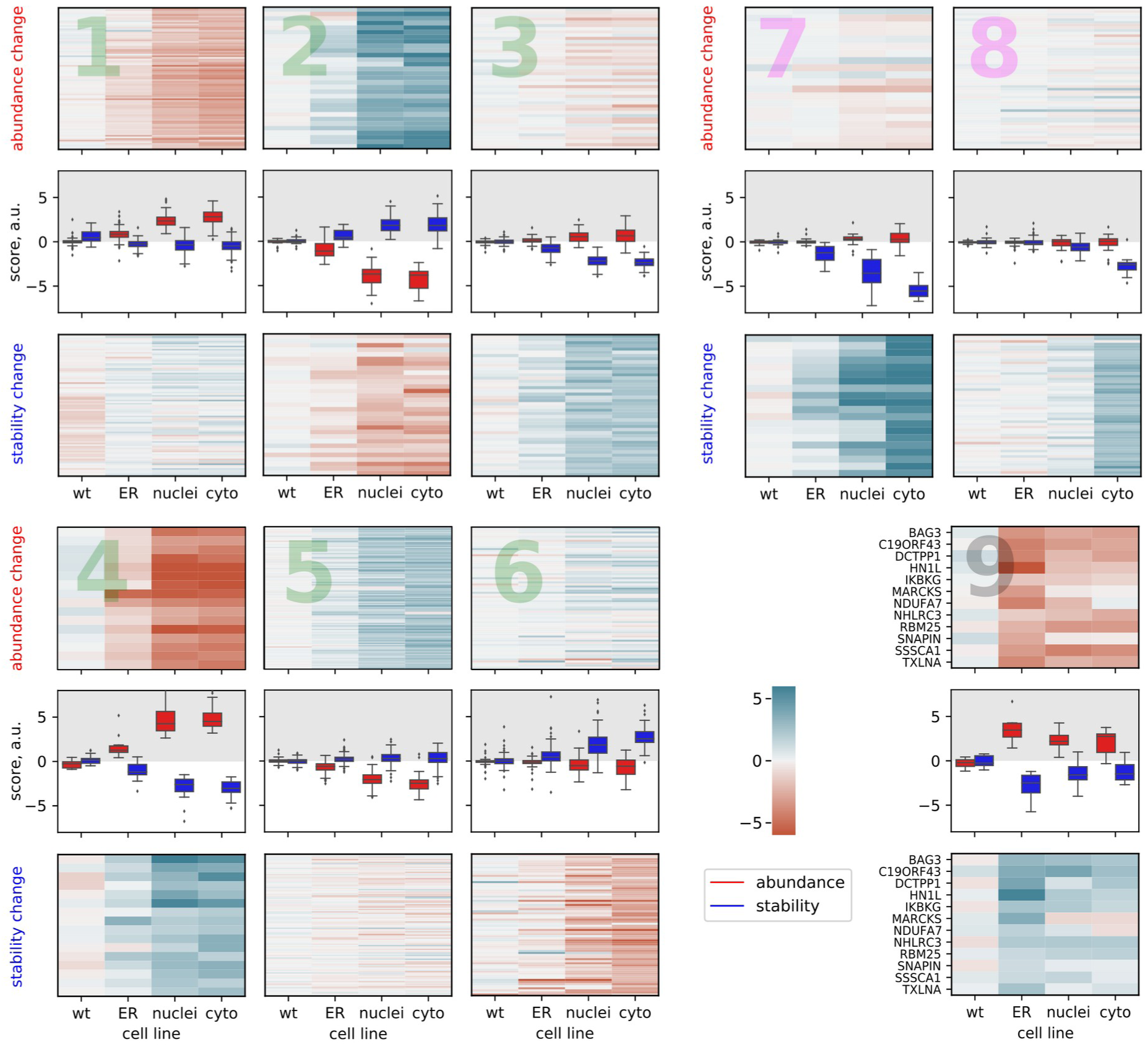
Protein groups with significantly different response profiles upon compartment-specific H_2_O_2_ generation. Panels demonstrate the results of k-means cluster analysis; the nine clusters were united into three groups: 1-6, 7-8, and 9. Clusters which belong to the same group are denoted by numbers of the same colour. Each cluster is represented by a vertical panel of three figures. The top and the bottom panes show heatmaps of protein abundance and stability scores (Figure S2B), respectively. The middle figure depicts distribution of abundance and stability scores of proteins for a particular stable cell line (N=3 biol. repl.).

### Identifying protein complexes involved in cellular response to H<u>_2_</u>O_2_ generation

Through clustering, hit proteins were grouped and analysed for ontology enrichment, providing more detailed insights than would be gained from analyzing the entire set of hit proteins (Figure 3A). Gene ontology enrichment analysis (GOEA) across three domains — cellular component, biological process, and molecular function —showed significant overrepresentation of proteins associated with protein synthesis, RNA-binding, and ribosome formation in cluster 1, which included many ribosomal proteins (over 40) such as 60S ribosomal proteins L6 (RPL6), L7a (RPL7A), and L13 (RPL13), as well as 40S ribosomal proteins S3a (RPS3A), S6 (RPS6), and S23 (RPS23) (Figure 3A). These proteins had high abundance scores, indicating an increased presence of these proteins in the soluble fraction upon H_2_O_2_ generation (Figure 2). This aligns with reports that protein synthesis (mRNA translation) is a crucial target of various cellular stressors, including H_2_O_2_-mediated oxidative stress^36,37^. Additionally, H_2_O_2_ can modulate mRNA translation at multiple levels, and due to inhibition of translation initiation, often leading to a shift from polysomes (multiple ribosomes actively translating a single mRNA) to monosomes^37,38^.

**Figure 3.**
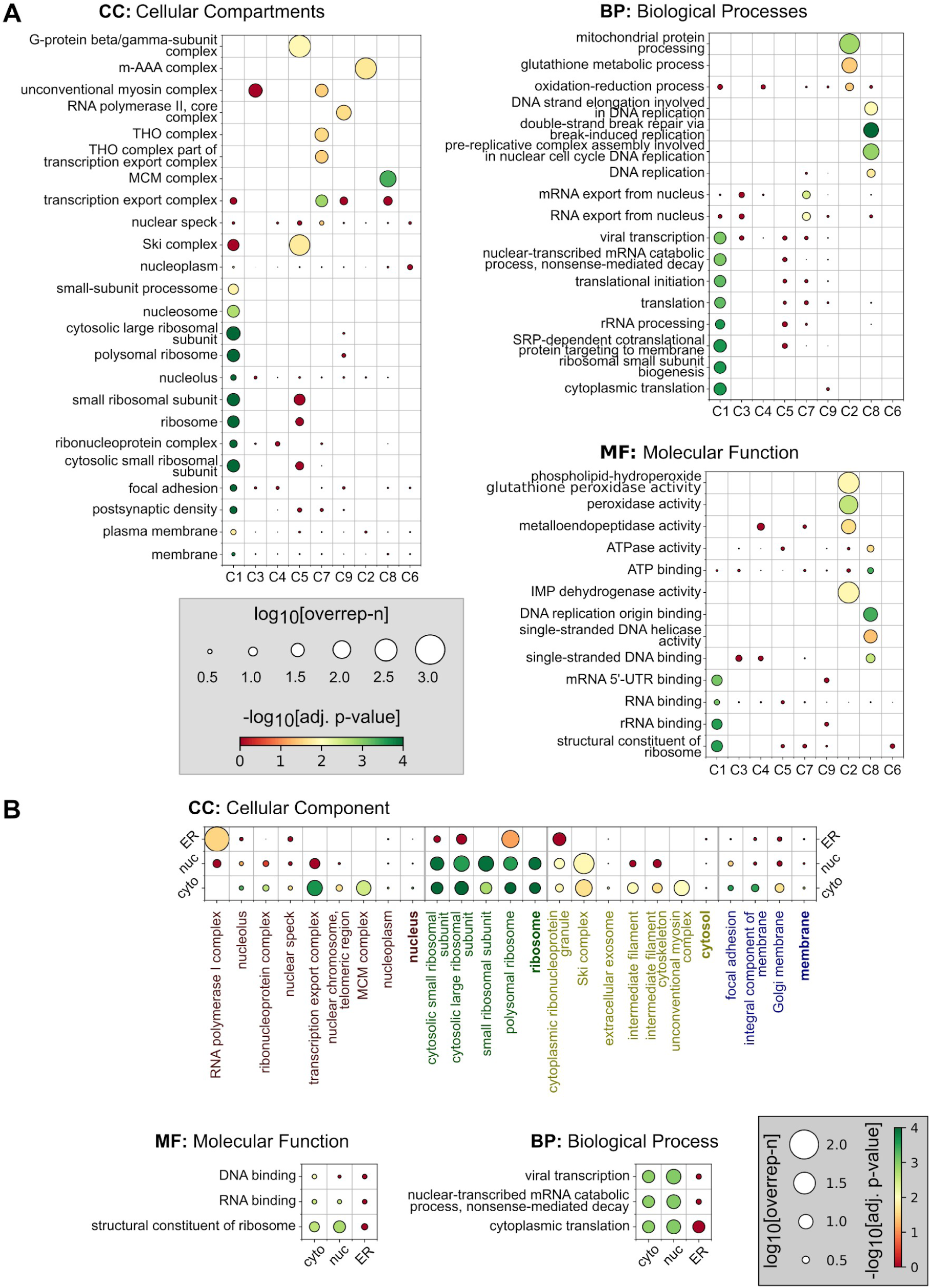
Results of gene ontology enrichment analysis performed for derived clusters (A) and for three cell lines stably expressing DAO (B). Colored frames separate clusters of different groups. Gene ontology terms were utilised from^63,64^. Adj. p-value – adjusted p-value, overrep-n – overrepresentation.

A unique feature of TPP is its ability to track entire protein complexes. Physically interacting proteins tend to co-aggregate upon heat treatment, resulting in similar melting curves^33^, which can be seen in our study, particularly with the helicase SKI2W (SKIV2L) and tetratricopeptide repeat protein 37 (TTC37) proteins of the SKI complex, which is enriched in cluster 4 (Figure 3A). The SKI protein complex represents a tetramer of SKIV2L, TTC37, and WDR61 (WD repeat-containing protein 61) with 1:1:2 stoichiometry, exhibits helicase activity (SKIV2L) and is necessary for the functioning of the exosome in the cell cytoplasm^39,40^. The exosome complex performs 3’-to-5’ mRNA digestion activity and participates in non-stop decay (NSD), a process that removes mRNA molecules with ribosomes stalled at the 3’-end of mRNAs due to poly(A)-tail readthrough^41^. Interestingly, H_2_O_2_ seemed to influence the solubility of the SKI complex at low temperatures, while the complex aggregated and precipitated rapidly at higher temperatures, independently of H_2_O_2_ (Figure S3A). There is evidence suggesting that the impact of H_2_O_2_ on the SKI complex, as yeast mutants of SKI complex components are highly sensitive to H_2_O_2_, is possibly due to the translation of aberrant mRNAs and the production of abnormal proteins^42^. As NSD was demonstrated to exist in mammalian cells^43^, it is plausible to suggest that the SKI complex may also play a role in eliminating NSD substrates that accumulate under oxidative stress in mammalian cells.

Another identified protein complex is the multi-subunit nuclear THO complex, consisting of six proteins (THOC1-3, 5-7), with THO complex subunits 1, 2, and 5 (THOC1, THOC2, THOC5) enriched in cluster 5 (Figure S3B). The THO complex is a core part of the larger and dynamic TREX (Transcription and Export) complex, linking gene transcription, mRNA processing, and mRNA export from the nucleus to the cytosol^44,45^. The processing of some mRNA molecules in the nucleus is coordinated by the THO complex via interaction with ZC3H14 (zinc finger CCCH-type containing 14)^46^, an RNA-binding protein involved in several post-transcriptional processes, such as regulation of length of the mRNA poly(A)-tail^47^, mRNA splicing^48^, and nuclear export^49^. Similar melting curves of ZC3H14 and components of the THO complex indicate their interaction (Figure S3B). While the THO and TREX complexes have been poorly studied in the context of oxidative stress, components like THOC1 and THOC5 play roles in cellular responses to oxidative stress^50,51^. For example, one study showed that astrocyte-conditioned media supported neuronal viability via PI3K-signaling under a glutamate-induced oxidative challenge^50^ via THOC1 and ZC3H14.

The core RNA polymerase II (Pol II) complex, with its largest proteins RPB1, RPB2, and RPB3 (POLR2A, POLR2B, POLR2C), was also identified upon H_2_O_2_ generation with high stability scores (cluster 6) (Figure S3C). Pol II participates in generating several RNA species, particularly mRNA^52^. One study showed that HeLa S3 cell nuclear extracts treated with 10 mM H_2_O_2_ for 15 min showed no transcriptional activity on an undamaged DNA template, indicating possible transcription repression via protein oxidation^53^. Another study reported that treating HeLa cells with 0.3 mM H_2_O_2_ for 10 min resulted in global Pol II promoter-proximal pausing, concluded to be due to suppressed aberrant transcriptional termination and unaffected initiation^54^, leading to higher promoter and enhancer occupation by Pol II and respective depletion of its soluble form. Thus, stabilisation of DNA-bound paused Pol II at promoters and enhancers upon H_2_O_2_ treatment may underlie the observed increase in thermostability of Pol II components. This assumption is consistent with results from a study by the Savitski group, showing that DNA-binding increases Pol II thermostability^34^.

Clusters 7 and 8 combined proteins with high absolute stability but not abundance scores. Cluster 7 was enriched in proteins with metallo-endopeptidase activity, including matrix ATPases associated with diverse cellular activities like the AAA (m-AAA) proteases paraplegin (SPG7) and AFG3-like protein 2 (AFG3L2); and the intermembrane AAA (iAAA) protease ATP-dependent zinc metalloprotease YME1L1 (YME1L1). These proteins are involved in mitochondrial protein quality control^55^.

Cluster 7 also included peroxidases like glutathione peroxidase 1 (GPX1), phospholipid hydroperoxide glutathione peroxidase (GPX4), and peroxiredoxin-5 (PRDX5), known for their roles in oxidative stress response and redox processes. Cluster 8 was enriched in proteins involved in the organization of the mini-chromosome maintenance protein complex (MCM), which functions as a heterohexameric helicase in DNA replication^56^. The MCM complex consists of six replication licensing factors (MCM2-7), five of which (MCM3-7) had high stability scores in the cyto-DAO cell line (Figure S3D). Although replicating DNA is sensitive to oxidation, the mechanisms connecting replication and cellular redox homeostasis are largely unexplored. The MCM3 subunit interacts with KEAP1, a redox-sensitive regulator of NRF2 degradation^57^. However, whether KEAP1-MCM3 interaction has an impact on regulation of DNA replication under oxidative stress conditions remains unclear^57,58^. Nevertheless, given the multiple PTMs that proteins of MCM complex can acquire^59^, and the ability of oxidative stress to affect PTMs (e.g., due to inactivation of protein phosphatases^60–62^), other mechanisms independent of KEAP1 may underlie the altered thermostability of the MCM complex. Other clusters were not significantly enriched in any GO terms.

To assess overrepresented GO terms in the three cell lines stably expressing DAO, another GO enrichment analysis (GOEA) was performed (Figure 3B). The nuc-DAO and cyto-DAO cell lines displayed considerable overlap, sharing many GO terms, with more terms overrepresented in the cyto-DAO cell line. This is unsurprising, as most hit proteins responded similarly to H₂O₂ in the nuc-DAO and cyto-DAO cell lines (Figure 2). Both cell lines were enriched in GO terms related to protein synthesis, RNA-binding, and ribosome formation (cluster 1). In contrast, the ER-DAO cell line exhibited distinct GO term enrichment (Figure 3B). While most proteins showed high abundance and stability scores in the nuc-DAO and cyto-DAO lines but lower scores in the ER-DAO, RNA polymerase I subunits RPA1, RPA2, and RPA49 (POLR1A, POLR1B, POLR1E) were significantly overrepresented only in the ER-DAO cell line.

### The influence of cytosolic H_2_O_2_ generation on protein interactions

Proteins can change thermostability for reasons other than altered protein-protein interactions^28^. Therefore, we selected four hits from the TPP experiment for a detailed study: PARK7 (cluster 6), TRAP1 (cluster 8), MAP2K1 (cluster 8), and UBA2 (cluster 6).

PARK7, a multifunctional protein, has a wide range of reported activities including deglycase^65^, protease^66^, and chaperone functions, which are particularly important under oxidative stress^67^. TRAP1 (tumor necrosis factor receptor-associated protein 1) is a mitochondrial HSP90-like chaperone (HSP75) involved in regulating protein synthesis, protecting from ER stress^68^, preventing mitochondrial permeability transition pore opening, and regulating mitochondrial respiration^69^. MAP2K1 (MEK1) is a key component of the RAF/MEK/ERK (MAPK) signaling pathway. Lastly, UBA2 is a subunit of an E1-activating enzyme involved in protein SUMOylation^70^.

FLAG-tagged versions of these proteins were expressed in HEK293 cells, with or without the induction of cytosolic H_2_O_2_ via the addition of 2 mM D-Ala. These FLAG-tagged proteins were then used as bait in a co-immunoprecipitation (co-IP) experiment, followed by the quantification of protein interactors using LC-MS/MS (Figure S4A). This approach, known as affinity-purification mass-spectrometry (AP-MS), was carried out under five conditions: CPD, CP−, C−D, −PD, and −P−. Here, C/− indicates the cyto-DAO or wild-type (control) cell lines, P/− signifies the presence or absence of expression of a FLAG-tagged protein of interest (POI) in cells, and D/− denotes the presence or absence of 2 mM D-Ala in the cell medium (Figure S4B). The treatment conditions in the AP-MS experiment were designed to closely mirror those used in the TPP procedure (Figure S4A). After co-IP, the FLAG-tagged POI levels were assessed using SDS-PAGE and Western Blotting (WB) (Figure S5).

The quality of the co-IP procedure was assessed by quantifying the amounts of the protein of interest (POI) and HyPer-DAO in each sample (Figure S6A). As expected, FLAG-tagged POIs were not expressed in the C−D condition, resulting in low levels of their endogenous counterparts. Variation in POI quantities between conditions with and without POI expression depended on the specific POI, likely reflecting differences in POI overexpression levels and the efficiency of magnetic bead washing and protein elution. For UBA2, the difference between the C−D and other conditions was minimal, possibly due to lower UBA2-FLAG overexpression compared to other POIs (Figure S5). HyPer-DAO was only expressed in the cyto-DAO cell line, and thus was expected to be present solely in the CPD, CP−, and C−D conditions, not in the −PD or −P− conditions. However, trace amounts of HyPer-DAO were detected in all samples, including those from the wild-type (wt) cell line. This may be due to incomplete washing of the HPLC column in the LC-MS/MS setup, leading to carryover of peptides from previous cyto-DAO samples into wt samples.

To gain a deeper understanding of the protein composition in the derived samples and their similarities, we directly compared the results obtained for the same POI under different conditions. As shown in Figure 6B, each sample contained at least several hundred identified proteins, with most being shared across all conditions for the specific POI. Interestingly, the number of proteins identified in POI-expressing samples was comparable to those in mock-transfected conditions (i.e., C−D or POI-free). This suggests that a large portion of proteins in the samples were likely non-specific binders to the magnetic beads, regardless of POI presence and despite multiple washing steps in the co-IP procedure (Figure 6B, no expression control).

Next, we conducted an AP-MS experiment to identify proteins whose interactions with selected POIs change in response to cytosolic H_2_O_2_ generation, focusing on three groups of proteins (Figure 4A). The first group (Ox) included proteins whose levels were altered in response to H_2_O_2_ (red circles). The second group (Bait) comprised proteins whose levels changed in response to POI expression in cells (blue circles). The third group (Ctrl) contained proteins that altered their levels in response to both POI expression and H_2_O_2_ generation (grey circles). Proteins from the Ox group were considered non-specific binders and were excluded from further analysis after statistical evaluation (Figure 4B). Examples of these proteins include Myotubularin-related protein 14 (MTMR14), MOB kinase activator 2 (MOB2), and heat shock 70 kDa protein 1A (HSPA1A). Their increased binding to beads upon H_2_O_2_ treatment could be due to H_2_O_2_-induced conformational changes or post-translational modifications that enhance binding. Alternatively, H_2_O_2_ may directly promote interactions between these proteins and multiple binding partners.

**Figure 4.**
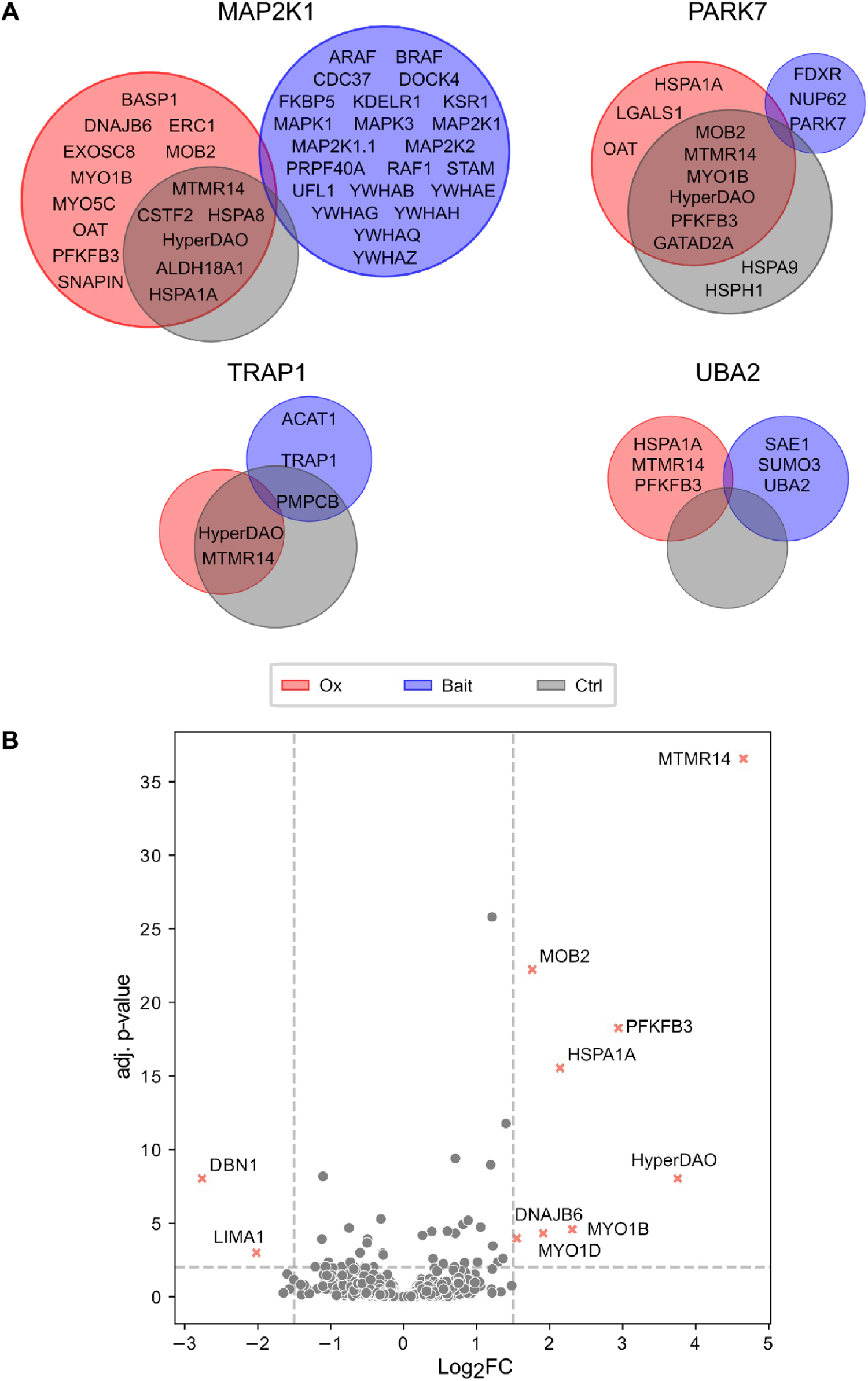
Proteins identified in AP-MS experiments with MAP2K1, TRAP1, UBA2, and PARK7 baits. (A) Differences in the interactomes of four proteins selected for analysis. Groups of proteins: “Ox” – CPD and C−D vs CP−, WPD and WP− conditions; “Bait” – CPD, CP−, −PD and −P− vs CD; “Ctrl” – CPD vs CP−, −PD and −P−, where C/W in the first position stands for the cyto-DAO or wt (control) cell lines, P/− – presence/absence of expression of a FLAG-tagged protein of interest (POI) in cells, and D/− – presence/absence of a D-Ala (2 mM) in cell medium. (B) Distribution of proteins identified in samples of the AP-MS experiment, according to the ratio (Fold Change) of the mean protein amount in samples derived from cells where H_2_O_2_ generation was induced (CPD and C−D conditions) to the mean amount in samples derived from cells where it was not (CP−, −PD and −P− conditions), with an indication of the respective statistical significance. Adj. p-value – adjusted p-value. N = 3 biol.repl.

### TRAP1

We identified two proteins whose interaction with TRAP1 was significantly altered by oxidative stress: β-MPP (PMPCB) and ACAT1 (Figure 4A). PMPCB, along with PMPCA, forms the mitochondrial processing peptidase (MPP), which is a part of the mitochondrial protein import machinery and removes N-terminal mitochondrial targeting sequences (MTS) from preproteins^71^. While changes in PMPCA levels were not statistically significant, the expression pattern closely matched that of PMPCB (Figure S7). Our results indicate that the presence of PMPCA and PMPCB correlates with TRAP1, suggesting a direct or indirect interaction between TRAP1 and the MPP complex. Furthermore, both PMPCA and PMPCB were found in significantly higher levels in samples with cytosolic H_2_O_2_ generation, implying that H_2_O_2_ may enhance TRAP1-MPP binding, highlighting a potential role for this interaction under oxidative stress conditions (Figure S7).

ACAT1, acetyl-CoA acetyltransferase 1, is another mitochondrial enzyme that catalyses the formation of acetoacetyl-CoA from two acetyl-CoA molecules. It shares several common interactors with TRAP1, as revealed by systematic AP-MS experiments performed with 21 mitochondrial bait-proteins^72^. Our findings, supported by the STRING database protein network^73^, show a close relationship among PMPCA, PMPCB, ACAT1, and TRAP1 (Figure 5). We also identified other TRAP1-associated proteins, such as SUGT1, HADHA, HSPD1, and UQCRC2, in TRAP1 samples. However, their levels did not show the same correlation with TRAP1 as observed with PMPCA, PMPCB, and ACAT1 (Figure S7).

**Figure 5.**
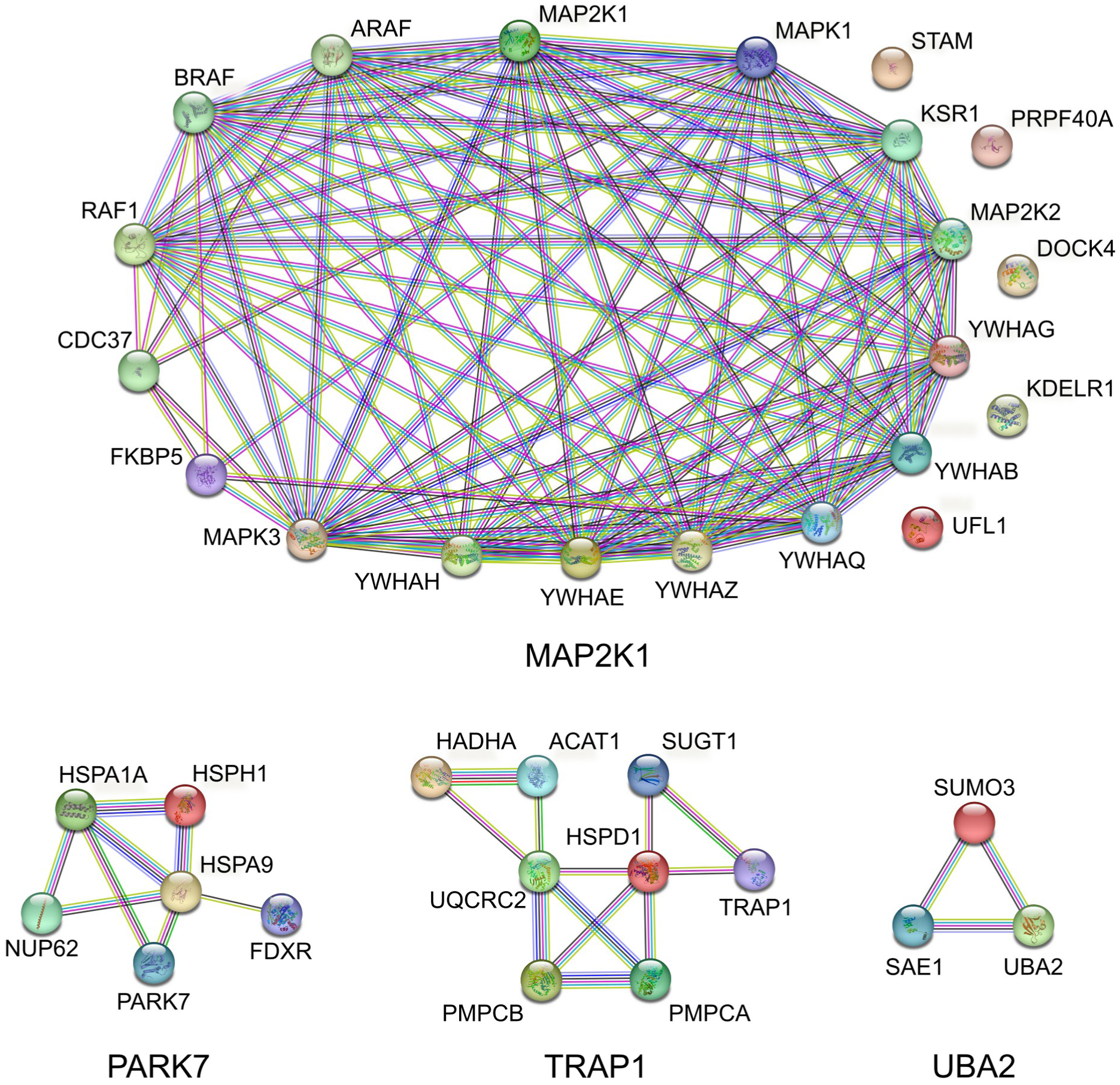
Interaction networks between proteins of interest and their partners identified in our AP-MS experiments. We included proteins, whose presence was found to be significantly affected in co-immunoprecipitation experiments, as well as with some other (related) proteins according to the STRING database^73^.

### PARK7

In samples where PARK7 was used as a bait protein and cytosolic H_2_O_2_ was generated, several proteins, including heat shock protein 105 kDa (HSPH1) and mitochondrial HSP70 – HSPA9 (GRP/mtHSP70/mortalin), were enriched, suggesting an increased interaction between PARK7 to these proteins in response to H_2_O_2_ (Figures 4A and S7). The literature strongly supports the role of PARK7 in oxidative stress. For instance, one study reported an association and colocalization of HSPA9 with PARK7 in HEK293T cells, significantly enhanced by treatment with 100 µM H_2_O_2_ for 4 h^74^. Similarly, another study found a low level of interaction between PARK7 and HSPA9 under normal conditions, which substantially increased in resveratrol-pretreated H9c2 cardiomyocytes undergoing hypoxia-reperfusion, accompanied by mitochondrial translocation of PARK7^75^. Moreover, the oxidation-induced formation of C106-SO_2_H (cysteine sulfenic acid) was shown to lead to mitochondrial localization of PARK7 and increased cell viability, a phenomenon not observed with the C106A PARK7 mutant^76^. Finally, PARK7 was identified as a component of the IP_3_R-GRP75-VDAC complex, which contributes to the formation of functional mitochondria-ER contacts (MERCs) and the transport of calcium ions between these two organelles^77^. Given the well-established interplay between ROS and calcium, including the ability of calcium to modulate mitochondrial ROS production, it is tempting to speculate that PARK7 might also regulate calcium transport between these organelles under increased ROS conditions, thus further affecting mitochondrial ROS generation^78^.

Although HSPA1A was previously identified as a “non-specific binder,” the significant increase in HSPA1A levels in PARK7-overexpressing samples compared to controls suggests a strengthened interaction with PARK7 upon H_2_O_2_ exposure. This aligns with a study showing that HSPA1A enhanced its interaction with FLAG-tagged PARK7 in a co-IP experiment in the HCT116 cells under TNFSF10-mediated oxidative stress^79^.

Apart from these two chaperones, the levels of ferredoxin reductase (FXDR) and nucleoporin 62 (NUP62) also correlated with PARK7’s presence in samples (Figure S7). According to the STRING database, both proteins can interact with PARK7 via HSPA9 (Figure 5). However, unlike HSPA9, H_2_O_2_ generation did not affect their levels in PARK7-overexpressing samples (Figures 4B and S7). Overall, our AP-MS results suggest that H_2_O_2_-triggered changes in PARK7’s thermostability can be attributed to an altered interactions with chaperone proteins. Nevertheless, given that cysteine 106 of PARK7 is known to be redox-sensitive^80^, oxidative modifications of this cysteine may also play a role.

### MAP2K1

The RAF/MEK/ERK (MAPK) signaling pathway is a well-known cascade of protein kinases that regulates various cellular processes^81^. MEK, a kinase in the middle tier of this pathway, is activated by RAF kinase and subsequently activates ERK. MEK functions as a stable heterodimer consisting of MEK1 (MAP2K1) and MEK2 (MAP2K2), which, despite their high sequence similarity, have distinct roles^82^. Previous studies using MAP2K1 as bait in co-IP experiments have identified several interactors, including upstream kinases (ARAF, BRAF, RAF1), downstream kinases (ERK2 (MAPK1), ERK1 (MAPK3)), and adapter proteins (KSR1 and various 14-3-3 proteins)^82,83^. In our study, we identified additional proteins in samples overexpressing MAP2K1 that were absent in mock-transfected controls. According to the STRING database, some of these proteins, such as CDC37 and FKBP5, belong to the MAP2K1-centered protein network. However, H_2_O_2_ did not appear to affect the binding of these proteins to MAP2K1. Thus, the high absolute stability score of MAP2K1 observed in our TPP experiment likely stems from factors other than altered protein interactions, such as post-translational modifications.

### UBA2

Protein SUMOylation is a post-translational modification mediated by the covalent attachment of Small Ubiquitin-like Modifier (SUMO) proteins to target proteins. This process, similar to ubiquitination, involves several key enzymes: E1 activating enzyme (a heterodimer of SAE1 and UBA2), E2 conjugating enzyme UBC9 (UBE2I), and one of several E3 ligases^70^. SUMOylation is reversible, with SUMO proteases (isopeptidases) removing SUMO proteins from their targets. Some components of this SUMOylation machinery are regulated by ROS and H_2_O_2_. For example, the catalytic cysteine residues in the E1 activating (UBA2) and E2 conjugating (UBC9) enzymes can form intermolecular disulfide bonds upon oxidation, leading to enzyme inactivation^84^. Isopeptidases such as SENP1, 2, and 3 are also known to be redox-regulated^70^.

In our AP-MS experiment using UBA2-FLAG as bait, several known UBA2 interactors were identified compared to non-transfected controls. Among these interactors were SAE1, which forms a heterodimer with UBA2, and SUMO2/3 (two isoforms that are 95% identical and typically indistinguishable), which were present at much higher levels than SUMO1, reflecting their known expression differences^85^. Notably, the levels of SAE1 and SUMO2/3 proteins did not vary between conditions with or without H₂O₂, indicating that their interactions with UBA2 are not affected by H_2_O_2_. Consequently, understanding the high stability score of UBA2 determined in our TPP experiment likely arises from mechanisms that warrant further investigation.

## Discussion

Redox processes play a fundamental role in living systems, yet many proteins involved remain unidentified. In this study, we aimed to uncover proteins driving the cellular response to H_2_O_2_ generation within distinct compartments. We engineered stable HEK293 cell lines expressing HyPer/TagBFP-DAO proteins in the cytosol, nucleus, or ER lumen and applied a two-dimensional thermal proteome profiling (2D-TPP) approach.^35^ DAO specifically activates localized H_2_O_2_ production in response to D-ala, leading to targeted disruptions in the redox balance. This controlled generation of H_2_O_2_ puts strain on scavenging enzymes like catalase, glutathione peroxidases, and peroxiredoxins, which rely on reducing equivalents such as NADPH and glutathione to detoxify ROS. The increased demand for these reducing agents can deplete their levels, diverting them from other essential metabolic processes and weakening overall redox homeostasis.

Using 2D-TPP, we analysed the response of over 5000 proteins to compartment-specific H_2_O_2_ production. We identified several hundred proteins with significant changes in abundance or thermostability, providing a comprehensive catalogue of proteins responsive to compartment-specific H_2_O_2_ production, both downstream of and independent of thiol oxidation. Our findings emphasize the cell’s vulnerability to localized oxidative stress and the critical role of reducing equivalents in maintaining redox balance and defending against oxidative damage.

Under conditions of intracellular H_2_O_2_ production, we expected to identify proteins directly reacting with H_2_O_2_ or involved in redox relays. This was confirmed by the detection of thioredoxin reductases TXNRD1 and TXNRD2, key components of the thioredoxin antioxidant system^86^, as stability hits across all three engineered cell lines^87,88^. Additionally, we identified peroxiredoxins (PRDXs), except for PRDX1. Notably, PRDX5 was stabilized in the cytosolic and nuclear cell lines, whereas PRDX4 showed no stability changes in any condition.

We also detected strong thermal stabilization of SUMO E1-activating enzyme, SAE1 and UBA2, consistent with previous TPP results from HepG2 cells exposed to exogenous H_2_O_2_^13^. Another significant hit was ubiquitin carboxyl-terminal hydrolase 5 (USP5), a deubiquitinating enzyme, which exhibited reduced stability in the cyto-DAO cell line.

Cluster and gene ontology analysis revealed that the cellular responses to H_2_O_2_ generated in the cytosol and the nucleus were highly similar but distinct from the ER-DAO cell line. The largest number of hits occurred in the cytosol, followed by the nucleus, with far fewer hits in the ER-DAO cell line, potentially due to differential DAO expression or the ER’s oxidizing environment, which limits further oxidation ^11^. The cytosol and nucleus showed slight differences, possibly due to variations in DAO expression levels and the more reducing nuclear environment. However, the similarities between these two compartments suggest limited diffusion of H_2_O_2_ or migration of proteins between these compartments.

Our TPP analysis revealed enrichment of various protein complexes, including ribosome subunits, RNA polymerase, and the MCM complex. Ribosomal proteins, in particular, were more abundant, possibly due to a shift from polysomes to monosomes, a response previously linked to H_2_O_2_-induced attenuation of global protein synthesis^89^. Given that oxidative stress can lead to mutations in mRNA, reducing protein synthesis may be a protective mechanism to prevent the production of faulty proteins^38^. While the role of ribosomes in oxidative stress adaptation is established, other complexes like the THO complex require further study.

In our AP-MS experiments with MAP2K1, PARK7, TRAP1, and UBA2, we observed significant changes in interactomes for TRAP1 and PARK7. While the interactions of PARK7 with identified chaperones under oxidative stress conditions are relatively well understood, TRAP1’s interaction with the mitochondrial processing peptidase (MPP) complex is less explored. The role of TRAP1 as a chaperone and MPP’s role in mitochondrial protein import suggest this interaction could be related to impaired protein import during oxidative stress^90^, though further experiments are needed to confirm this.

It is important to note the substantial difference between AP-MS and TPP experiments. AP-MS uses FLAG-tagged bait proteins expressed at higher levels, while TPP examines endogenous protein levels. Therefore, the absence of altered protein-protein interactions for MAP2K1 and UBA2 in AP-MS doesn’t necessarily rule out the possibility of changes in thermostability due to protein-protein interactions. Furthermore, in the case of UBA2, the lack of UBC9 detection, which forms a disulfide bond with UBA2 under oxidative stress, prevents its use as a control for assessing the impact of H_2_O_2_ on the UBA2 interactome. Beyond changes in protein-protein interactions, the altered thermostability of MAP2K1 and UBA2 could also be attributed to post-translational modifications (PTMs) or other factors.

In conclusion, our study sheds light on the complex and multifaceted nature of the cellular response to H_2_O_2_, highlighting the involvement of proteins critical to a wide range of cellular functions—from H_2_O_2_ detoxification to transcription, translation, and replication. Notably, this response is highly dependent on the cellular compartment where H_2_O_2_ is generated. These findings pave the way for future research, enhancing our understanding of the intricate dynamics of cellular redox biology.

## Methods

### Plasmid Assembly

The pLVX-Puro plasmid backbone required for the assembly of the pLVX-Puro-HyPer-DAO-3NLS and pLVX-Puro-HyPer-DAO-NES plasmids was amplified by PCR from the pLVX-Puro-DAAD plasmid using the 8-2 and 6-7 pairs of primers (Table 2). Parts of the HyPer-DAO-NES and HyPer-DAO-3NLS fragments were amplified using the 9-11 and 9-10 pairs of primers from the AAV-HyPer-DAO-3NLS plasmid. The amplicons were assembled into target plasmids following a previously described procedure^91^. To generate the pLVX-Puro-TagBFP-DAO-KDEL plasmid, pC1-TagBFP-DAO-KDEL was amplified using the 86-87 pair of primers and the obtained fragment was digested with the XbaI/BamHI (NEB, R0145/R0136) pair of endonucleases along with pLVX-Puro-DAAD and then ligated. All source plasmids were taken from V.Belousov’s lab.

**Table 1:** Plasmids.

| plasmid | origin |
| --- | --- |
| AAV-HyPer-DAO-3NLS | V. Belousov’s lab |
| pC1-TagBFP-DAO-KDEL | V. Belousov’s lab |
| pLVX-Puro-DAAD | V. Belousov’s lab |
| pMD2.G | Addgene, #12259 |
| pCMV-dR8.91 | Creative Biogene, OVT2971 |
| pcDNA3.1(+)-PARK7-FLAG | GenScript, NM_007262.5 |
| pcDNA3.1(+)-TRAP1-FLAG | GenScript, NM_016292.3 |
| pcDNA3.1(+)-MAP2K1-FLAG | GenScript, NM_002755.4 |
| pcDNA3.1(+)-UBA2-FLAG | GenScript, NM_005499.3 |
| pcDNA3.1(+) | Invitrogen, V79020 |

**Table 2:**
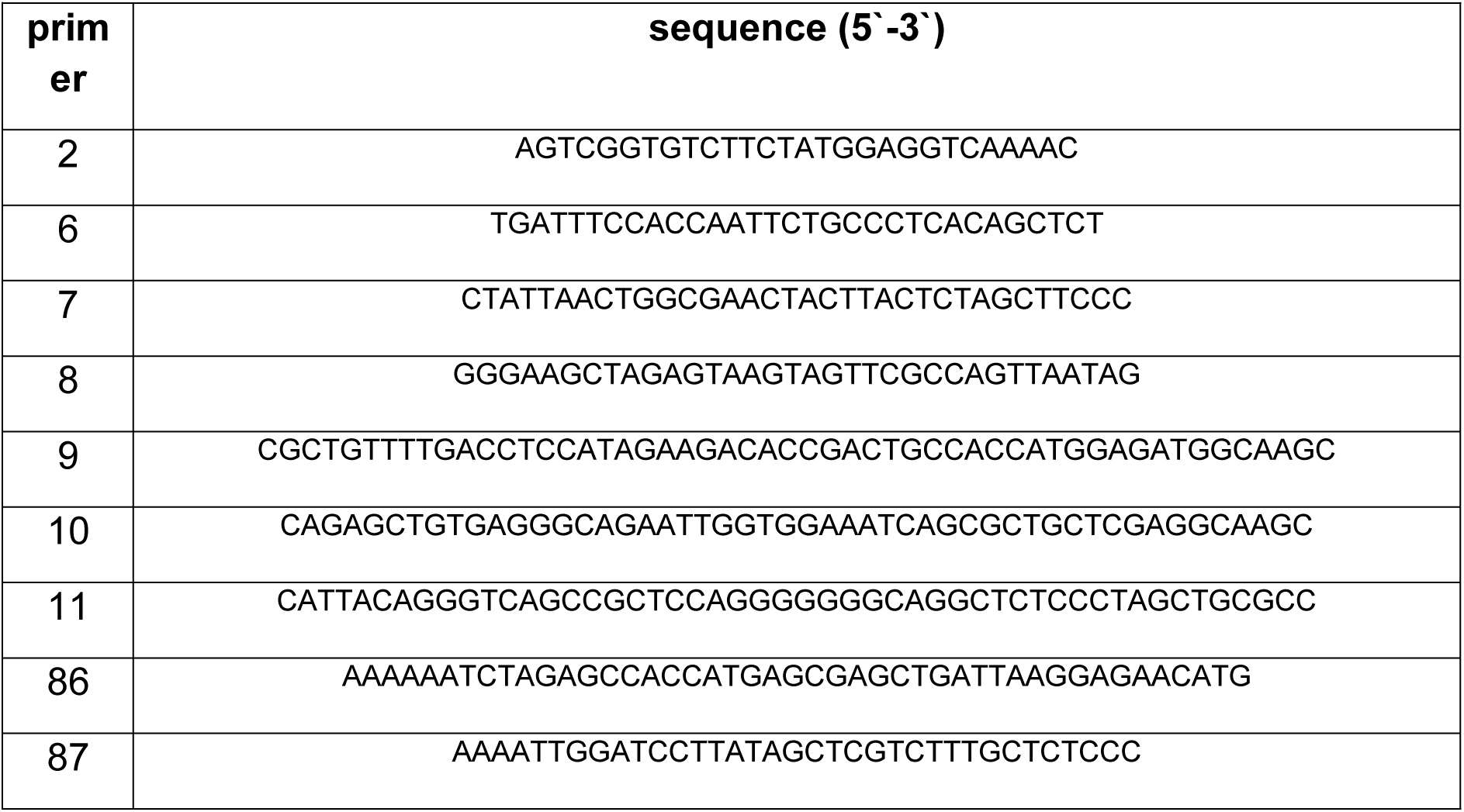
Primers.

### Cell Culture

All procedures with eukaryotic cell lines were fulfilled in sterile conditions of a biosafety cabinet. The original (wt) HEK293 cell line and its genetically modified derivatives were grown in DMEM-medium (Thermo/Gibco, 41966-029) with FBS (10%) at 37 °C and CO_2_ (5%). Cells were cultured in DMEM containing Pen/Strep (1%) only prior to and during the AP-MS experiment. The original HEK293 cells, HEK293 cell lines stably expressing DAO (produced in the current study) and HEK293TN cells were regularly split 1:3, 1:2-3, 1:10 every 3-4 days, respectively using trypsin solution. All cell lines were regularly checked for mycoplasma contamination according to the manufacturer recommendations (PCR Mycoplasma Test Kit I/C, PromoKine, PK-CA91).

### Stable Cell Line Production

The puromycin (VWR, 540222-25) concentration required for selection of stable HEK293 cells in 48 h was determined in advance as 1 µg/ml. All procedures involving the virus were carried out in an S2 biosafety level lab. To produce stable HEK293 cell lines, HEK293TN cells were seeded in DMEM (10% FCS), and on the next day transfected with a mix of pCMV-dR8.91 (Creative Biogene, OVT2971), pMD2.G (Addgene, #12259) and pLVX-Puro (encoding a gene of interest) plasmids using FuGENE-HD (Promega, E2312). 24 h after transfection, the virus-containing medium was replaced with a fresh one and HEK293 cells were seeded for infection. 48 h after transfection, the viral-containing medium was filtered (0.45 µm), and 16 µg/ml polybrene was added to the filtrate. This medium was then used to replace the medium the HEK293 were growing in, and subsequently diluted 2x with fresh medium. On the next day the viral-containing medium was replaced with fresh medium. Starting from the next day (i.e. 48 h post-transduction), puromycin was added to the medium. Cells were incubated with puromycin for 2-3 days. Before transferring cells out of the S2 lab, cells were washed with PBS to remove remaining virus and virus-negative cells were further transferred to S1 conditions. HEK293 cell lines expressing HyPer-DAO-NES and HyPer-DAO-3NLS were used for monoclonal cell line production. Cells were seeded on 10 cm Petri dishes in a volume of 10 ml at a concentration of 5 cells/ml. Once they became discernible, single cells were marked as potential candidates, and dishes were incubated for several weeks. Separate colonies consisting of hundreds of cells were isolated using cloning cylinders, expanded and frozen.

### Fluorescent Microscopy

Prior to imaging, the stable HEK293 cell lines were incubated in Experiment Medium (see EM, Table 3) for 45 min at 37°C without CO_2_. Imaging was performed using a Zeiss Observer D1 microscope equipped with a Zeiss EC-Plan Neofluar 40x/1.3 Oil Ph3 objective (for tracing the oxidation kinetics of HyPer) and a Zeiss EC Plan Neofluar 10x/0.3 Ph1 (for stable cell line image acquisition), a Zeiss Axiocam 702 mono camera and a Colibri LED light source at 37°C. Cells expressing HyPer were imaged by the sequential excitation of cells with light of two wavelengths – 420 and 505 nm via 420/40 nm and 500/15 nm band-pass excitation filters, respectively, together with a 515 nm dichroic mirror and collection of emitted light via a 539/25 nm (emission) filter. Imaging of TagBFP was performed using 400 nm wavelength light and usage of another filter cube (band-pass excitation filter: 400/40 nm, dichroic mirror 458 nm, band-pass emission filter: 483/32 nm). The oxidation kinetics were analyzed using Fiji (ImageJ). The image background was subtracted using a rolling ball algorithm and the threshold was adjusted. The stack of images corresponding to 505 nm was divided frame-by-frame by 400 nm stack to obtain the HyPer ratio signal (R).

**Table 3:** Recipes of complete media and solutions.

| **medium/solution** | **recipe** |
| --- | --- |
| CM<br>(culture medium) | DMEM, high glucose, pyruvate,<br>FCS – 10%,<br>(Pen/Strep – 1%) |
| FLAG-peptide elution buffer<br>(5 mg/ml, 10x stock) | FLAG-peptide – 4 mg,<br>1x TBS – 0.8 ml |
| EM (Experiment<br>Medium) | DPBS calcium, magnesium, 1x,<br>Glucose – 5.56 mM |
| TPP lysis buffer | DPBS, 1x,<br>NP-40, 1.33%, |
|  | MgCl<sub>2</sub>, 2.5 mM,<br>cOmplete, 1.6x,<br>PhosStop, 1.6x,<br>Benzonase, 0.417 U/μl |
| AP-MS Lysis Buffer 1 | DPBS, 1x,<br>NP-40, 0.8%,<br>MgCl<sub>2</sub>, 1.5 mM,<br>cOmplete, 1x,<br>PhosStop 1x,<br>Benzonase, 0.250 U/μl |
| AP-MS Lysis Buffer 3 | DPBS, 1x,<br>NP-40, 0.8%,<br>MgCl<sub>2</sub>, 1.5 mM |
| AP-MS Lysis Buffer 4 | DPBS, 1x,<br>NP-40, 0.8%,<br>MgCl<sub>2</sub>, 1.5 mM,<br>cOmplete, 1x,<br>PhosStop 1x |

**Table 4:** Reagents, media, antibodies, kits.

| reagent | manufacturer | catalogue number |
| --- | --- | --- |
| Carboxy-terminal FLAG-BAP™ Fusion Protein | Sigma | P7457 |
| cOmplete™ Protease Inhibitor Cocktail | Sigma | 11697498001 |
| Puromycin dihydrochloride | VWR | 540222-25 |
| D-alanine | Sigma | A7377 |
| FLAG-peptide | Sigma | F3290 |
| FuGENE® HD Transfection Reagent | Promega | E2312 |
| XbaI | NEB | R0145 |
| BamHI | NEB | R0136 |
| PhosStop (phosphatase inhibitor) | Sigma | 4906845001 |
| Polyethylenimine, Linear, MW 25000, Transfection Grade (PEI 25K™) | Polysciences | 23966 |
| TMTpro™ 16plex Label Reagent Set | Thermo Scientific | A44522 |
| *Antibodies* |  |  |
| Monoclonal ANTI-FLAG® M2 antibody, mouse (primary) | Sigma | F3165<br>working dilution: 1:1000 |
| Goat anti-Mouse IgG (H+L) Secondary Antibody, HRP | Thermo Scientific, Invitrogen | 62-6520<br>working dilution: 1:5000 |
| *Kits* |  |  |
| ANTI-FLAG M2 Magnetic Beads | Sigma | M8823-5ML |
| Pierce™ ECL Western Blotting Substrate | Thermo Scientific | 32109 |
| PCR Mycoplasma Test Kit I/C | PromoKine | PK-CA91 |
| *Media* |  |  |
| DMEM, high glucose, pyruvate | Thermo Scientific/Gibco | 41966-029 |
| Opti-MEM™ (Reduced Serum Medium) | Thermo Scientific/Gibco | 31985062 |
| DPBS calcium, magnesium | Thermo Scientific/Gibco | 14040 |

D-Alanine on-the-fly addition was performed in two steps: first, 0.1 ml of 10 times more concentrated D-Ala solution in imaging medium was prepared (using 1M D-Ala water stock), second, this 0.1 ml was added to 0.9 ml of imaging medium in the cell dish.

### Thermal Proteome Profiling (TPP)

#### TPP Experimental Procedure

HEK293 cells were seeded into 15 cm dishes in 30 ml culture medium (DMEM/FCS), to obtain 60-70% confluency on the day of experiment. On the day of experiment the medium was changed to 15 ml Experiment Medium (EM, 1x DPBS with Mg^2+^/Ca^2+^/glucose) and dishes were incubated for 45 min at 37 °C without CO_2_. The EM was then replaced with either 15 ml EM supplemented with 2 mM (wt/nuc-DAO/cyto-DAO) or 8 mM (ER-DAO) D-Ala or EM without D-Ala and the cells were incubated for a further 10 min in the same conditions. Medium from each dish was collected to an individual 50 ml tube, and cells were treated with 3 ml of trypsin solution (+/– D-Ala) for 3 min at 37 °C. Trypsinization was stopped with 7 ml of CM (culture medium) (+/– D-Ala), and cells were transferred to the same 50 ml tubes. Dishes were rinsed with additional 10 ml of EM (+/– D-Ala), and the remaining cells were collected to the same 50 ml tubes. Cells were counted, centrifuged and resuspended in EM (+/– D-Ala) to have 5 x10^6^ cells/ml.

Ten aliquots of each cell suspension (0.1 ml) were transferred into a 0.25 ml 96-well PCR-plate. Plates were centrifuged for 2 min at 390 *g*, RT. A fraction of supernatant volume (80 µl) from each well was removed and cell pellets were resuspended in the remaining volume of supernatant. Plates were sealed with aluminium foil and subjected to a temperature gradient (ten temperature points: 37.0, 40.4, 44.0, 46.9, 49.8, 52.9, 56.5, 58.6, 62.0, 66.3℃) using a thermal cycler for 3 min. Plates were then incubated for an additional 3 min at RT.

All following lysis steps were performed on ice or at low temperature. Cold TPP lysis buffer (see Materials, 30 µl) was added to each well of a plate, and the content of each well was resuspended. The plate was sealed and incubated on a shaker for 1 h at 4 °C and with constant gentle stirring. Wells of a 96-well filter plate (0.45 µm) were pre-wet with lysis buffer (50 µl). The filter plate was centrifuged for 2 min at 250 g to dry the filters. The plate with samples was also centrifuged for 3 min at 700 g at 4 °C. The supernatant from the samples (40 µl) was transferred to wells of the filter plate, the filter plate was centrifuged for 5 min at 4°C and 500 x *g*, and the filtered supernatant was collected in a standard 96-well plate.

#### Mass Spectrometry (MS)

The protein concentration of samples exposed to the two lowest temperatures (37.0℃ and 40.4℃) was determined using a BCA assay kit, and the sample volumes containing 10 µg of protein were used for MS analysis. The volume of the samples exposed to the other temperatures used for MS analysis was equal to the mean volume of the two samples for which the concentration was determined.

The measurement procedure was performed according to the «LC-MS/MS measurement» section described in^34^ with minor modifications. Protein samples were treated similarly to the described procedure using a modified SP3 protocol^92,93^. Peptides were labelled with TMTpro (Thermo Scientific, A44522) (Figure S1B), and the labelled samples (four cell lines treated with or without D-Ala) corresponding to one pair of temperatures (16 samples in total) were mixed and fractionated, resulting in twelve fractions^92^. Peptides were separated using the same LC system and settings as described before^34^. The LC system was connected to the same Q Exactive™ Plus mass spectrometer (Thermo Scientific) and the mass spectrometry was performed according to the described procedure with the normalised collision energy equal to 30.

#### Data Analysis

Protein identification and quantification was performed using both the IsobarQuant^94^ and Mascot search engines using theTrEMBL human proteome as a reference database. Only proteins with at least two unique peptide matches in at least 2/3 of the replicates, were kept for the analysis. Batch effect was removed by the corresponding function of the LIMMA package. Finally, vsn normalization^95^ was applied to the log_2_ of raw quantity values. Normalisation coefficients were calculated for each temperature separately. Stability and abundance scores were computed as previously described according to the equations (Figure S2B)^93^. After calculation scores were normalised to Z(0,1) and fed to the lmFit LIMMA function. The LIMMA test was performed for the statistical estimation of stability and abundance Z-scores and for finding differences between +D-Ala and –D-Ala conditions for every cell line. Computed t-values were further evaluated with the fdrtool to obtain false discovery rates (q-values)^96^. In accordance with performed statistical analysis, proteins were classified as ‘hits’: ** (q-value < 1% and |Z-score| > 3), * (q-value < 5% and |Z-score| > 2), and ‘not hits’ (others). Hits were then clustered according to their behaviour under oxidative conditions. For every protein the following vector was built:

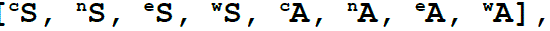

where the superscript indicates the cell line (c – cyto-DAO, n – nuc-DAO, ER-DAO, w – wt), S – stability score, A – abundance score. Then K-means clustering was performed and the optimal number of clusters (9) was determined with the elbow method.

#### Gene Ontology Enrichment Analysis

Gene ontology enrichment analysis (GOEA) was fulfilled using the Python goatools package^97^. Basic ontology (go-basic.obo) was used as an ontology graph from: Ashburner et al., 2000; Carbon et al., 2021^63,64^, and goa_human.gaf was used for mapping, respectively. The list of all identified proteins in TPP was used as background.

### Affinity Purification Mass Spectrometry (AP-MS)

#### PEI transfection

HEK293 cells were seeded in 15 cm dishes at 60% confluency in DMEM (10% FCS, Pen/Strep). On the next day, 1 ml OptiMEM (Thermo/Gibco, 31985062),

4.26 pmol of plasmid DNA (15-20 µg) and PEI 25K™ (Polysciences, 23966) (DNA:PEI = 1:5 w/w) were mixed by tapping, incubated for 10 min at RT and the mixture was added to the cells. After 4 h, the cell medium was replaced with fresh one. Cells were incubated for a further 24 h before starting experiments.

#### Co-immunoprecipitation (Co-IP)

Five 15 cm dishes with HEK293 cells (2 × wt, 3 × cyto-DAO) were seeded as described in the “PEI transfection section”. The next day, four of the five dishes (2 × wt and 2 × cyto-DAO) were transfected with pcDNA3.1 plasmid encoding one of FLAG-tagged proteins of interest (POIs) and one (cyto-DAO) with empty plasmid (control) using PEI. After 48 h the medium was removed, cells were washed with 10 ml of 1x DPBS, and prewarmed EM (no D-Ala) (15 ml) was added to cells. Dishes were incubated for 45 min at 37°C without CO_2_. EM was replaced with 15 ml of EM with 2 mM or without D-Ala. Dishes were incubated for a further 10 min at 37 °C without CO_2_.

The medium from each dish was transferred to an individual empty tube (50 ml). Attached cells were collected by trypsinisation (3 ml) with/without D-Ala for 3 min at 37°C, stopped by addition of CM (7 ml) with/without D-Ala. Medium with cells was transferred to the same 50 ml tubes containing EM. 1× DPBS (10 ml, with/without D-Ala) was added to dishes to collect the remaining cells. Tubes were centrifuged for 15 min at 4°C and 100*xg*, and the supernatant was thoroughly removed. AP-MS lysis buffer 1 (see Materials, 0.5 ml) was added to each cell pellet, and cells were resuspended by pipetting. Suspensions were incubated on an orbital rotator for 1 h at 4°C, and then lysates were clarified for 15 min at 4°C and 15000 *x g*. Resin for co-IP (AntiFlag M2 magnetic beads, Sigma, M8823) was prepared (100 µl of suspension/reaction) by removing the storage buffer and washing the beads twice with lysis buffer 3 (0.25 ml/wash).

Cell lysates were added to washed beads. We made negative control using AP-MS lysis buffer 4 (see Materials, 0.5 ml) and positive control using AP-MS lysis buffer 4 (see Materials, 0.5 ml) including 6 µg of BAP-FLAG fusion protein (Sigma, P7457). Tubes were incubated for 1 h at 4°C using an orbital rotator. Beads were collected, the supernatant was removed, and beads were washed 3 times with AP-MS lysis buffer 4 (see Materials, 0.35 ml/wash). During the last washing step beads were transferred to a new tube, collected, and the washing solution was removed. Proteins bound to beads were eluted with 0.1 ml FLAG-peptide elution buffer (Table 3). Samples were incubated by gentle shaking for 30 min at 4°C. After incubation beads were discarded, the eluates were collected and stored for long-term storage at –20°C.

#### Western Blotting (WB)

Western blotting was performed using primary mouse monoclonal ANTI-FLAG® M2 antibodies (Sigma, F3165) and secondary goat anti-mouse antibodies conjugated to HRP (Invitrogen, 62-6520). Working dilutions were 1:1000 and 1:5000, respectively. Protein bands were detected using Pierce™ ECL Western Blotting Substrate kit (Thermo, 32109) and images were acquired using ChemiDoc XRS+ System (BioRad).

#### Preparation for MS and MS/MS

Protein samples (eluates from co-immunoprecipitation step) were precipitated with acetone (Biosolve), resuspended in 50 mM NH_4_HCO_3_ (Sigma) by sonication and digested overnight with trypsin/Lys-C at 37°C (Promega, MS grade). The reaction was stopped with 0.1% (v/v) TFA (trifluoroacetic acid, Biosolve). The lysate were then processed and subjected to UHPLC-LC-MS/MS analysis according to the procedure as described previously with minor modifications^98^. Peptide peaks above the threshold in the MS survey were dynamically excluded for 40.0 s. MS scans were in the m/z range from 350 to 1800. Top N peptide precursors were subjected for MS/MS and fragmented in HCD using normalized collision energy (NCE) at 30%. Automatic gain control (AGC) was set to 4×10^5^ and a maximum injection time (MaxIT) to 100 ms.

#### Raw data processing and analysis

The Thermo RAW files were initially converted into the mzML format using the ThermoRawFileParser. Next, a database search was performed using MSFragger 3.5 with the following parameters: a true precursor tolerance of 12 ppm, a fragment mass tolerance of 300 mDa, and the allowance of up to 3 missed cleavages. Two types of modifications, namely methionine oxidation and N-terminal acetylation, were employed. The database used in the search was obtained by augmenting the proteome UP000000589 with standard contaminants from the Philosopher package and the HyPer-DAO-NES sequence. The results were statistically refined using peptideProphet (based on MZID files) and proteinProphet (based on PROT.XML file), and label-free quantification (LFQ) was carried out using IonQuant 1.8 with the match-between-run option enabled, generating TSV files as output.

### Subsequent data processing and analysis

TSV files derived as output on the previous step were further processed using the R programming language. First, each TSV file (dataframe) was filtered to keep only proteins with the maximum number of unique spectral counts (among all samples) > 2. Next, proteins were filtered based on the Razor Peptides columns (maximum of Razor Peptides values among all conditions for a particular protein > 1). Then, MAP2K1 sample −P− no.1 was considered artificial and excluded. Proteins were then filtered one more time based on (razor) intensity values via converting a dataframe to a summarised experiment object (SE) and applying filter_proteins function to the obtained SE with parameters: “condition” and thr=0 (or thr=1 for UBA2 protein) specified. Data were then median normalised and missing data were imputed using the «impute» function with fun = «man», shift=2.6, scale=0.25 parameters for TRAP1 and fun = «MinProb», q = 0.005 (for PARK7 and UBA2 or q =0.01 for MAP2K1) for three other proteins. For PARK7 additional normalisation was performed using the “normalize_vsn” function from the “DEP” package.

Further analysis was performed using the “limma” package (R). Linear models were fitted using the “lmFit” function. Next, contrast matrices were built using following contrasts: 1) Ox (CPD and C−D vs CP−, −PD and −P− samples), 2) Bait (CPD, CP−, −PD, −P− vs C−D), 3) Ctrl (CPD vs CP−, −PD, −P) and the “contrasts.fit” function was applied. Then, empirical Bayes smoothing to the standard errors was performed (“eBayes” function) and p-values were adjusted using the Benjamini Hochberg method by applying the “p.adjust” function.

#### Non-specific binder determination

To identify proteins binding non-specifically to the magnetic beads upon intracellular H_2_O_2_ generation, CSV files derived in the previous step, were concatenated into a single dataframe. Proteins, which were not present in two or more of the initial CSV files were filtered out. Data was normalised by removing the mean and scaling to the mean std of all samples. A t-test (CPD and C−D vs CP−, −PD and −P− samples) was performed for each particular protein, and a multiple comparison correction was applied (alpha=0.05, method=“fdr_bh“).

### Data visualisation

Data visualisation was performed using the Python programming language. Third-party packages such as matplotlib (https://pypi.org/project/matplotlib/) and seaborn (https://pypi.org/project/seaborn/) were used for plotting most of presented figures in the current study. Venn (https://pypi.org/project/venn/) was employed for plotting Venn diagrams.

### Data Availability

The LC-MS/MS raw data files, peptide-to-spectrum matches, and peptide intensities from in TPP experiment are deposited in the PRIDE Archive database of the EBI-EMBL under the identifier PXD055538. The LC-MS/MS raw data files, extracted peptides, and protein intensities as well as other related data derived in AP-MS experiment are deposited under the identifier PXD041407.

## Materials

## Author declaration

Authors declare no competing interests.

## Acknowledgements

The authors thank Isabelle Becher, Frank Stein, and Mikhail Savitski from the Proteomics Core Facilities and Genome Biology Unit at EMBL-Heidelberg for performing the 2D-TPP protocol using our transfected cells, for conducting LC-MS/MS measurements on the resulting samples, and for providing the initial data analysis (Supplementary Table 1).

The paper is based on the materials of doctoral thesis of Dr. Aram Revazian “Cellular Response to compartmentalized redox alterations”, which are publicly available in eDiss portal of SUB Göttingen under License CC-BY 4.0 (http://dx.doi.org/10.53846/goediss-9408).

The authors are also grateful to Andrea Paluschkiwitz (Molecular Physiology, Department of Cardiovascular Physiology, University Medical Center, Georg-August-University, Göttingen, Germany) for the help with cell culture, fluorescent microscopy and various other lab-related issues. Authors would also like to thank Khadija Wahni (Brussels Center for Redox Biology & Structural Biology Brussels, Vrije Universiteit Brussel, B-1050 Brussels, Belgium) for the versatile help with organisational issues.

This research was funded by the Russian Science Foundation (RSF), grant number 23-75-30023. Generation of cell lines was partially funded by the Center for Precision Genome Editing and Genetic Technologies for Biomedicine (Grant No. 075-15-2019-1789 to Pirogov Russian National Research Medical University). AR acknowledges support by the Global Education Program of Skolkovo Institute of Science and Technology. IB acknowledges the Deutsche Forschungsgemeinschaft (DFG) SFB1190 Project P17, CRC1027 Project C4 and IRTG1816.

## Author contributions

Conceptualization, JM, VB;

Methodology: AR, CG, AN, DE, TL;

Formal analysis: AR, AN;

Investigation: AR, DV;

Resources: IB, JM, VB;

Data Curation: AN;

Writing – Original Draft: AR;

Writing – Review & Editing: AN, DE, JM, VB;

Visualization: AR, AN;

Supervision: VB, CG, IB, DE, JM;

Project administration: VB, AN, CG, IB, JM;

Funding acquisition: IB, VB, JM.

## Supplementary Information

**Supplementary Table S1**. [tpp2d-limma.xlsx].

**Complete table of all identified proteins in TPP-2D experiments.** Values of abundance and stability changes (Figure S2B) computed for +D-Ala/-D-Ala contrast are provided together with FDR values indicating the significance of each shift.

**Figure S1.**
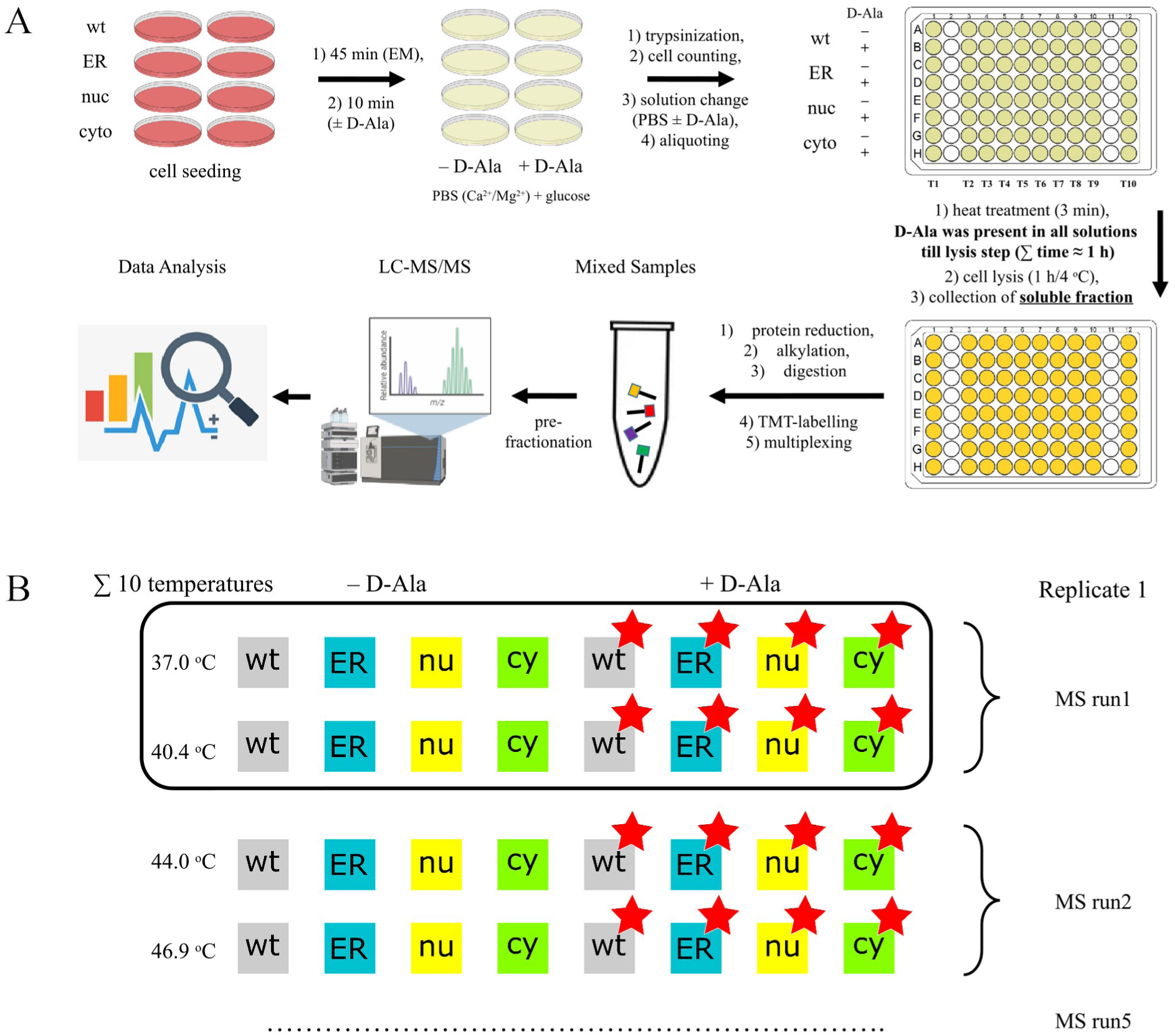
Visualisation of our 2D-TPP protocol including multiplexing strategy. **(A)** Workflow of Thermal Proteome Profiling experiment. Four HEK293 cell lines: wild type (wt), ER-DAO, nuc-DAO and cyto-DAO, were incubated with or without D-Ala. Next, cells were aliquoted and subjected to the thermal denaturation at different temperatures in a 37 – 65°C range and then lysed. The lysates were centrifuged and proteins from the soluble fraction (supernatant) were extracted and digested. Samples containing digested proteins were labelled according to the 2D-TPP scheme of multiplexing with 16-plex label reagent set TMTpro^TM^. Mixed labelled samples were prefractionated together, and analyzed using liquid chromatography with tandem mass spectrometry (LC-MS/MS) in a single MS run. **(B)** Scheme of samples multiplexing in MS. Samples derived from cells treated with D-Ala are labelled with red asterisks. Sixteen different samples corresponding to two temperatures were multiplexed in each MS run. Temperature range included ten different temperatures and thus five separate MS runs were required for one replicate. Note that each MS run is pre-fractionated, recorded, and then gathered at the data analysis step. Three biological replicates were implemented.

## Note S1. TPP experiment performance validation

To estimate the quality of the performed TPP procedure, we compared our identifications with the ones of three other recently performed TPP studies^1–3^. In the first study (Saei et al., 2020)^1^ authors performed TPP on intact HCT116 cells following a redox disturbance, using a thioredoxin reductase inhibitor, in the second one (Tan et al., 2018)^2^ authors employed a HEK293T cell line (close to HEK293 line, used in our study) for the experiment, while the last study (Becher et al., 2018)^3^ was performed using the experimental workflow and data processing pipeline similar to the ours. Total numbers of proteins in data sets of the compared studies were: 5010 (Saei et al., 2020), 7199 (Tan et al., 2018), 5391 (Becher et al., 2018) and 5252 (the current study) (Figure S2A). Among them, 3042 proteins were found to be common for all four data sets, which constitutes 57.9% of all proteins detected in the current study. Furthermore, the current data set shared 3761 (71.6%), 4400 (83.8%) and 3794 (72.7%) common proteins with (Saei et al., 2020), (Tan et al., 2018) and (Becher et al., 2018), respectively. Comparable number of proteins as well as significant share of common proteins between four studies clearly indicate a good proteome coverage in our study.

Saei A.A. et al. Comprehensive Chemical Proteomics for Target Deconvolution of the Redox Active Drug Auranofin. *Redox Biol*. **2020**, *32*, 101491. https://doi.org/10.1016/j.redox.2020.101491.

Tan C.S.H., et al. Thermal Proximity Coaggregation for System-Wide Profiling of Protein Complex Dynamics in Cells. *Science* **2018**, *359* (6380), 1170–1177. https://doi.org/10.1126/science.aan0346.

Becher I., et al. Pervasive Protein Thermal Stability Variation during the Cell Cycle. *Cell* **2018**, *173* (6), 1495-1507.e18. https://doi.org/10.1016/j.cell.2018.03.053.

**Figure S2.**
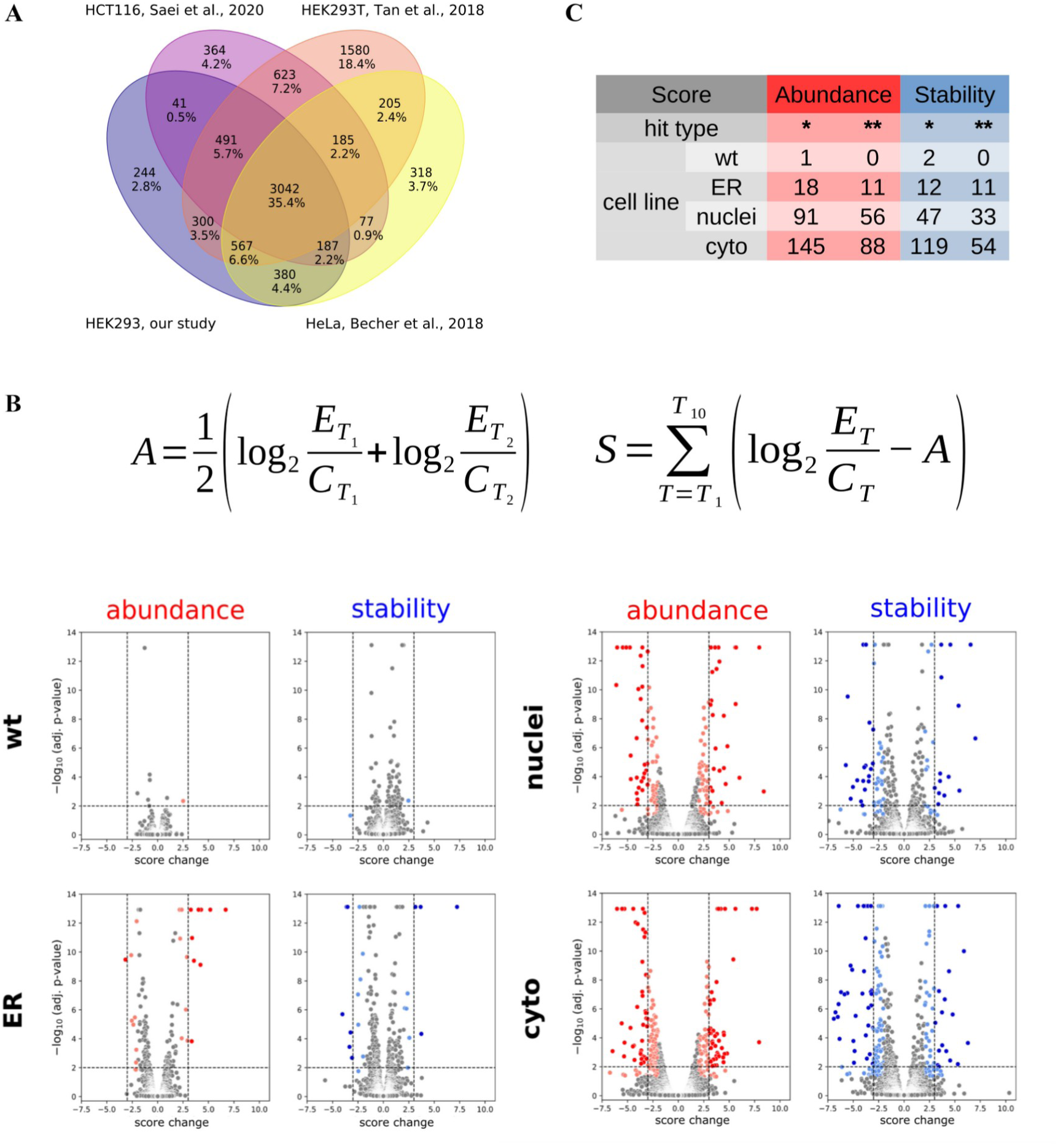
Initial analysis of 2D-TPP experiment. **(A)** Comparison of four indicated studies, wherein authors performed Thermal Proteome Profiling (TPP) on intact cells, based on sets of identified proteins (with at least two detected peptides each). Supplementary tables 4, 21 and 2 were used as data sets for comparison from (Saei et al., 2020), (Tan et al., 2018), and (Becher et al., 2018), respectively. Prior to comparison, supplementary table 21 (Tan et al., 2018) was filtered to remove proteins with less than two detected peptides. **(B)** Equations used for the calculation of protein abundance and stability scores. E_T_ and C_T_ – raw protein quantities corresponding to samples treated with (Experiment, E) or without (Control, C) D-Ala and heated at a particular temperature (T). T_1_ and T_2_ represent the first two temperatures of the range: 37.0 and 40.4 °C, respectively. **(C)** Numbers of proteins identified in TPP experiment in the current study with q-value < 0.01 and |Z-score| > 3 (**) and q-value < 0.05 and |Z-score| > 2 (*) in each of four HEK293 cell lines. **(D)** Distribution of hit (* – lighter and ** – darker colors, respectively) proteins in each of four HEK293 cell lines based on abundance/stability scores as well as statistical significance (N = 3 biol.repl.). Dashed lines indicate thresholds for hit (**) proteins.

**Figure S3.**
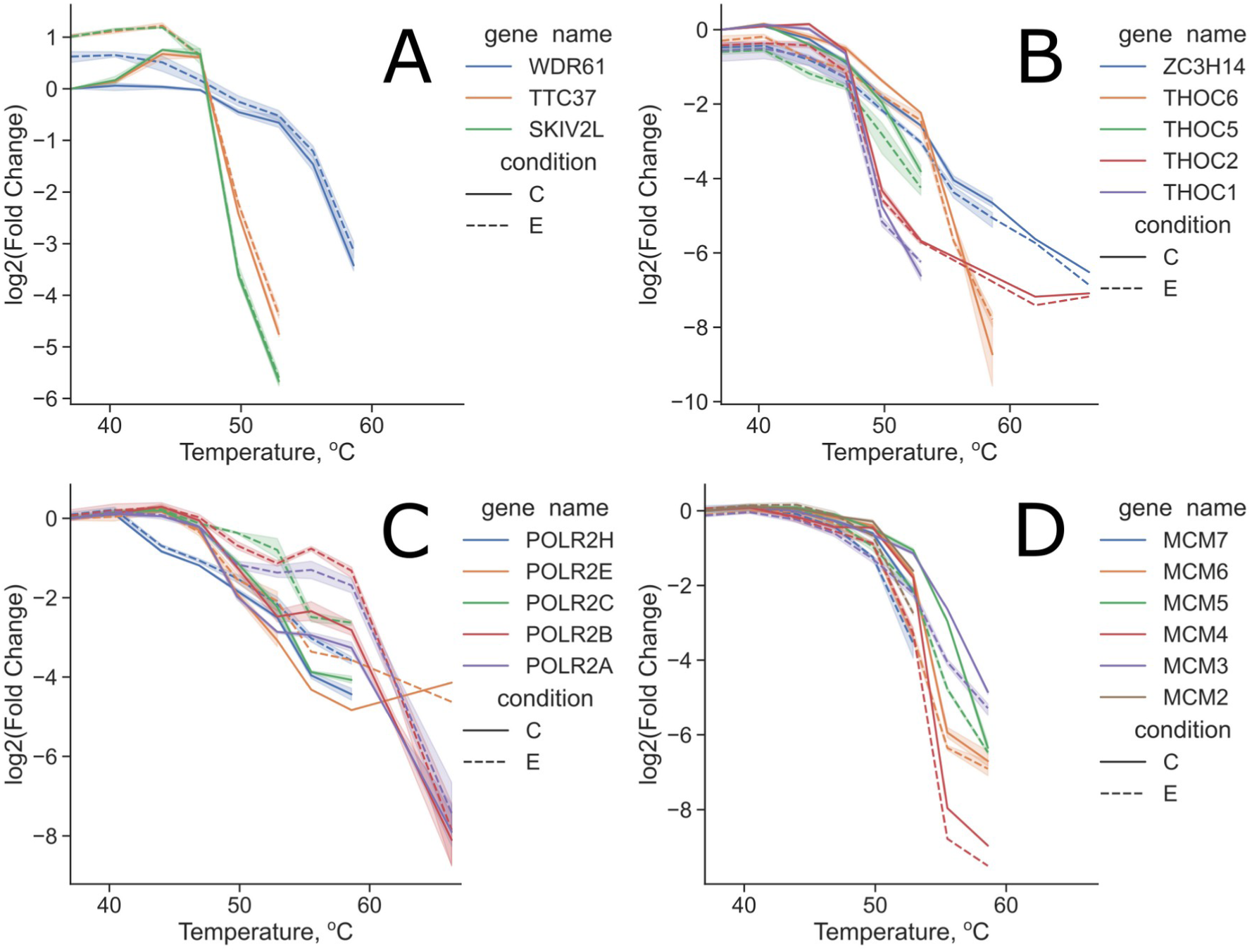
Melting curves of proteins participating in formation of certain protein complexes. (A) SKI complex, (B) THO complex, (C) RNA polymerase II and (D) MCM complex. Solid and dashed lines denote melting curves of proteins extracted from the cyto-DAO cell line treated with (experiment, E) or without (control, C) D-Ala. Protein amount at each temperature was normalized to the respective amount in the control at 37 °C (Fold Change). Shades around curves illustrate std (N = 3 biol.rep.).

**Figure S4.**
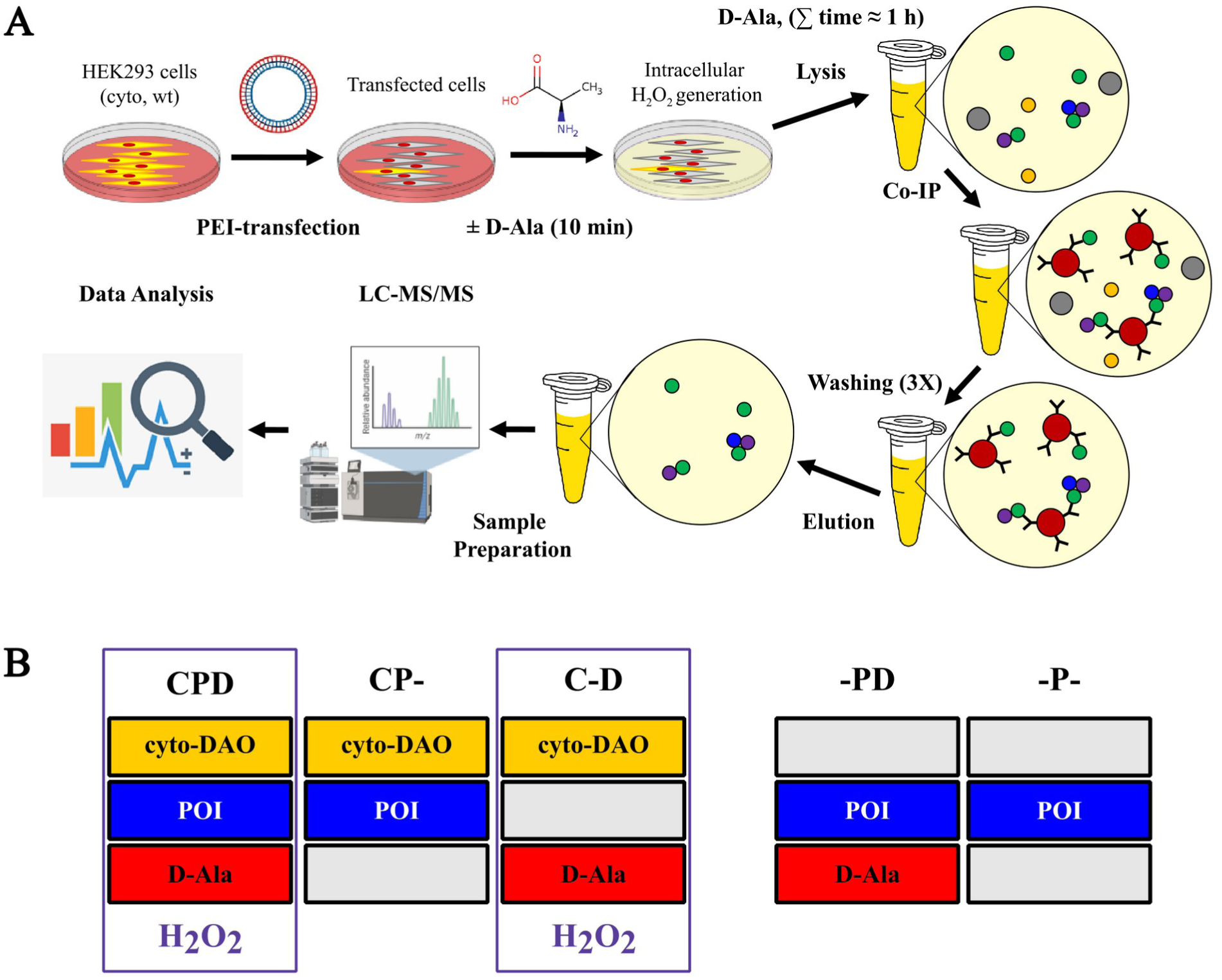
Working pipeline and experiment design of our AP-MS experiment. (A) Affinity purification – mass spectrometry (AP-MS) experiment workflow. (B) – Scheme of conditions used in co-immunoprecipitation (Co-IP) part of the AP-MS experiment. Each condition is denoted by a column of three rectangles. Top row rectangles indicate a HEK293 cell line that was used for the experiment: cyto-DAO (golden) or wild type (grey). Middle row rectangles depict a plasmid used for cell transfection: either a plasmid encoding FLAG-tagged protein of interest (POI, bait-protein) (blue) or an empty plasmid (grey). Bottom row rectangles display if D-Ala (2 mM) was added to cells. Presence of both golden (cyto-DAO) and red (D-Ala) rectangles in the column indicate H_2_O_2_ generation in cells.

**Figure S5.**
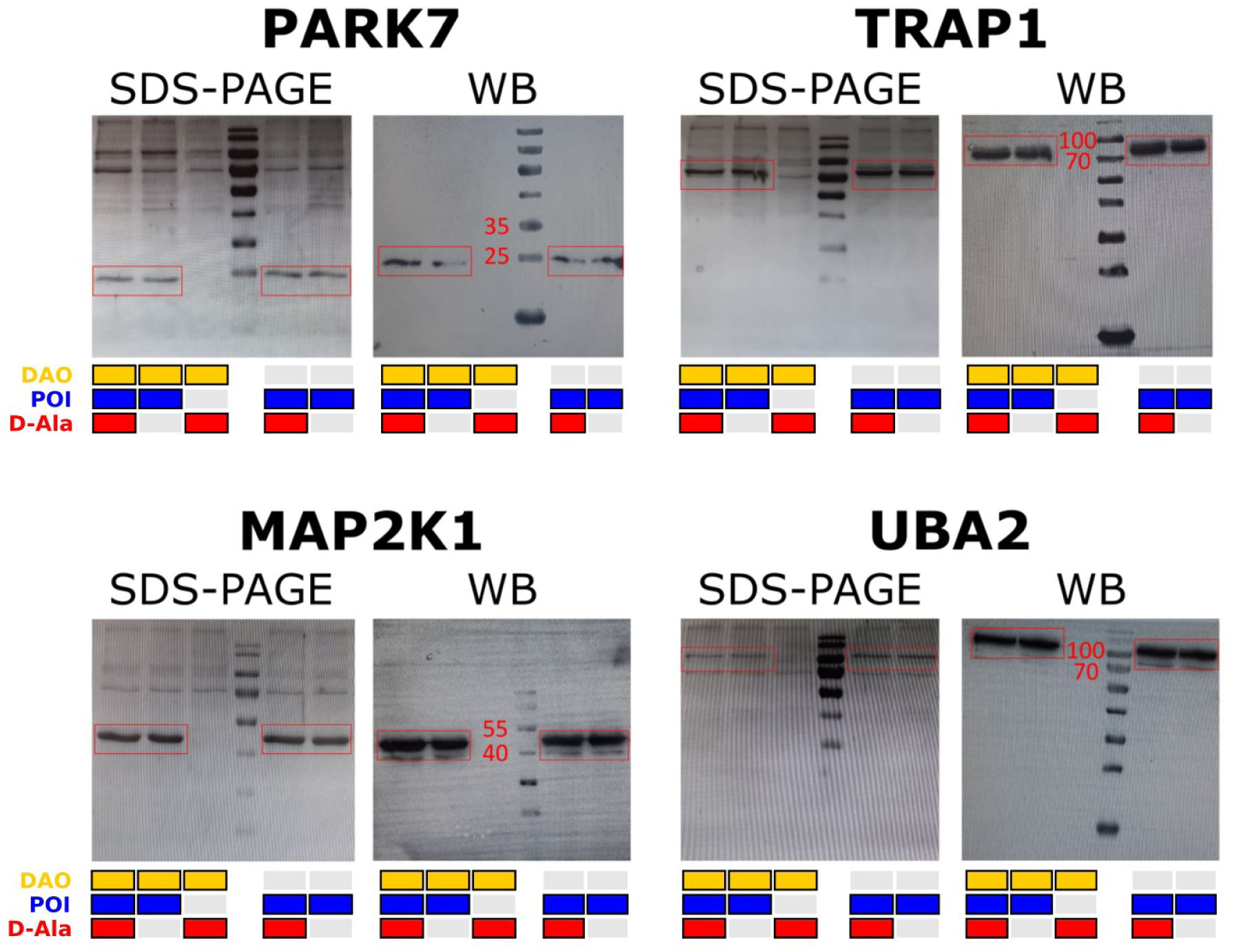
Examination of bait protein expression for AP-MS experiment. Validation of protein expression in HEK293 cells transfected with plasmids encoding one of four FLAG-tagged proteins of interest (POIs) or empty plasmid by SDS-PAGE and Western Blot (WB). After each SDS-PAGE proteins were transferred on a membrane, and a polyacrylamide gel with remaining proteins was stained with Coomassie Blue (SDS-PAGE). The membrane with transferred proteins was treated sequentially with primary (anti-FLAG) and secondary antibodies (WB). Molecular weights (observed) of target proteins: PARK7 – 20 (25) kDa, TRAP1 – 75 kDa, MAP2K1 – 49.2 kDa, UBA2 – 75 (∼100) kDa. Red frames display bands corresponding to POIs, red numbers – close molecular weights (kDa) of protein markers. Golden rectangles denote usage of the cyto-DAO cell line in the experiment (grey ones – wt cell line), blue ones – expression of a POI in cells and red ones – presence of D-Ala (2 mM) in cell medium. PageRuler™ Prestained Protein Ladder, 10 to 180 kDa (Thermo Scientific, 26617) was used as a marker.

**Figure S6.**
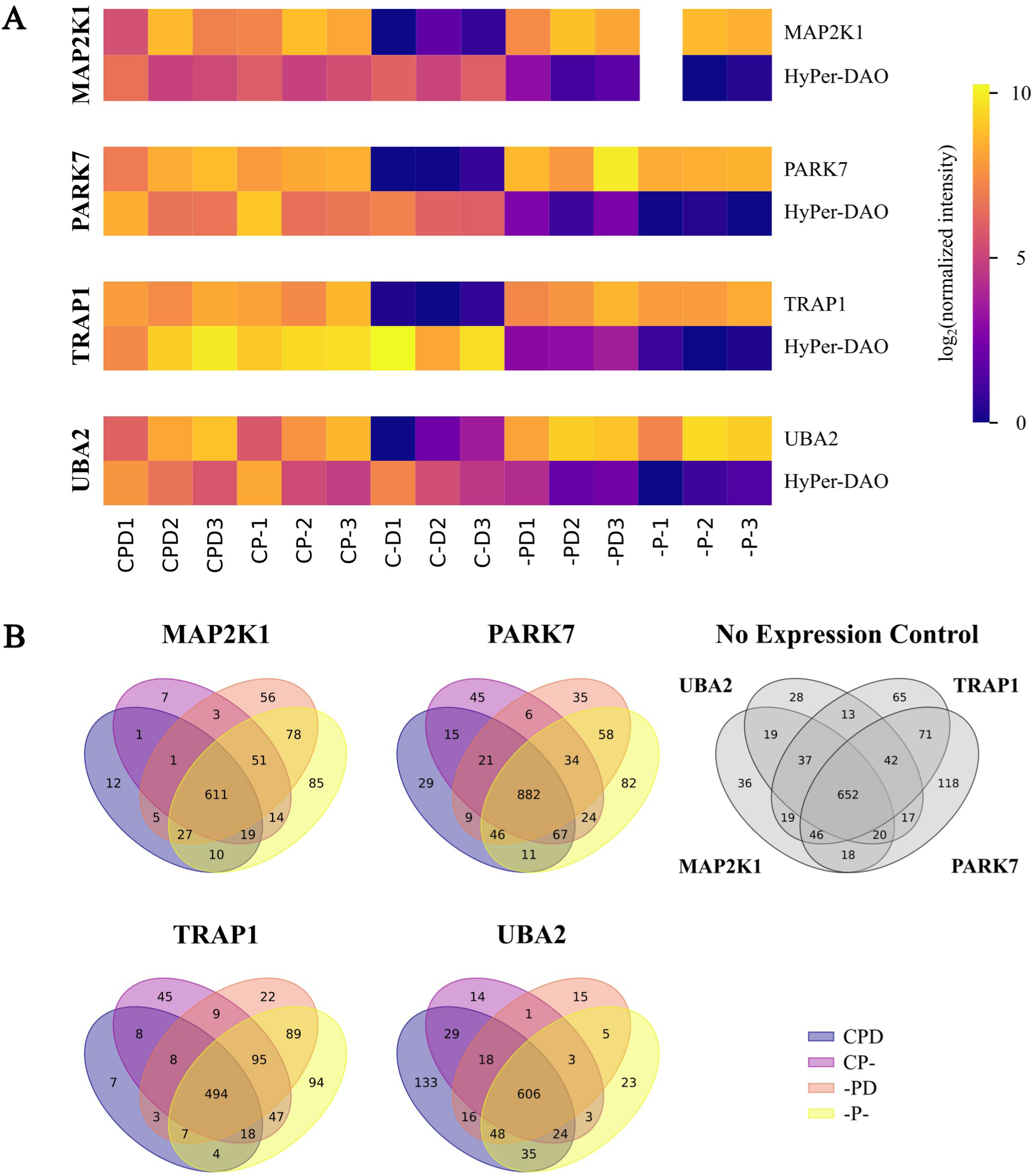
Quality evaluation of the AP-MS experiment. **(A)** Relative amounts (normalized to the minimum in a row) of proteins of interest (POIs) and HyPer-DAO proteins, identified in each sample for every experiment. The “−P−1” sample for the experiment with MAP2K1 was excluded from the data analysis and thus is missing. **(B)** Number of proteins identified in all samples corresponding to a particular condition, except for bait-protein free “C−D“ one (coloured Venn diagrams). “No expression control” diagram displays the number of proteins identified in samples corresponding to “C−D” conditions of experiments performed with different POIs (N = 3 biol.rep.). Coding of the experimental conditions are explained in Fig. S4B.

**Figure S7.**
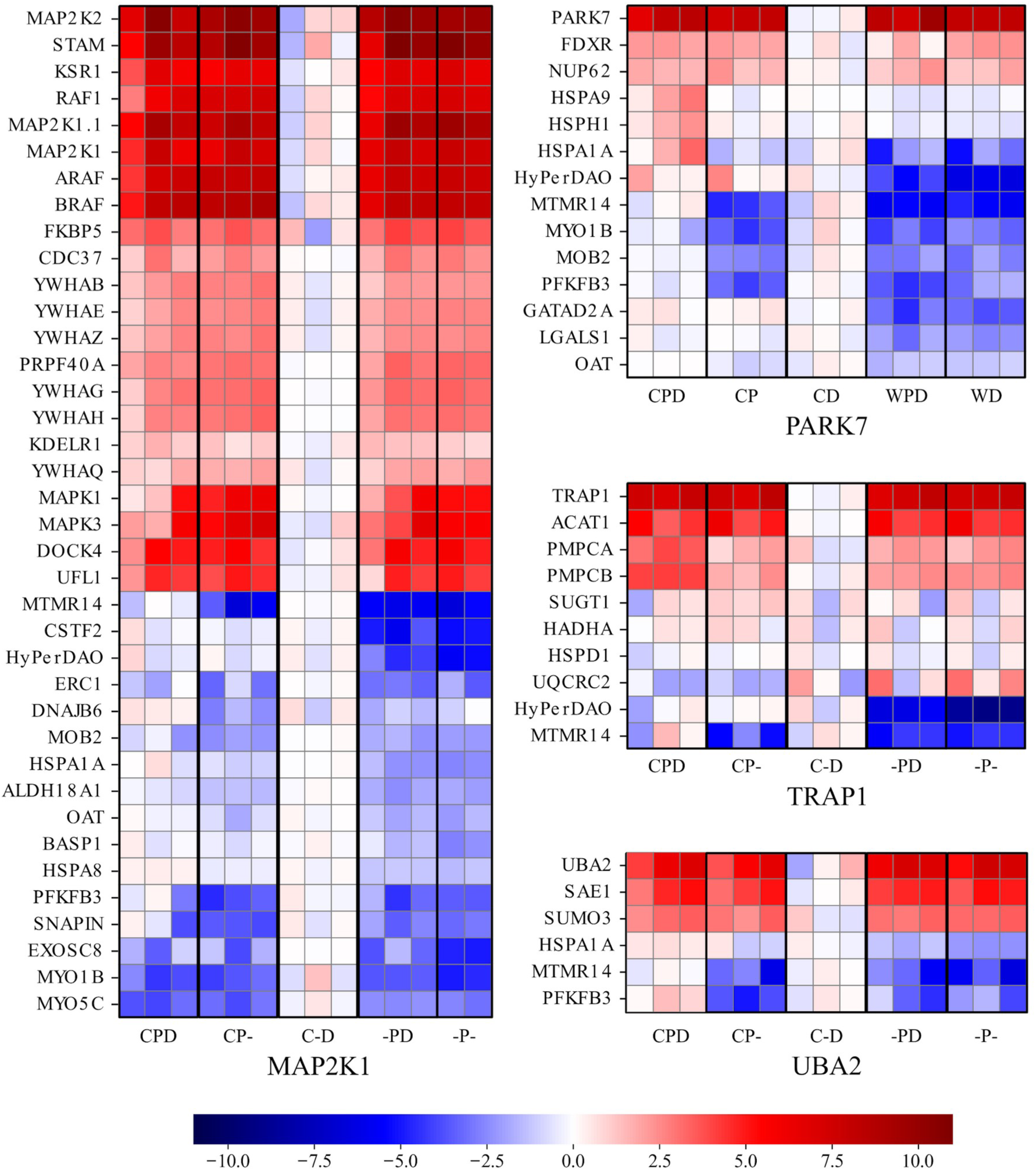
Heatmaps of all AP-MS experiment results, built for signifant protein set. On the heatmaps demonstrate for each sample the relative amount of proteins, which altered amounts depending on the existence of a FLAG-tagged bait-protein (protein of interest, POI) and depending on H_2_O_2_ generation in cells. Amounts of some proteins related to POIs according to the STRING database (Figure 5) and proteins, which were identified as “non-specific binders” (Figure 4B) are represented for comparison as well. Protein amounts in each row were normalized to the mean value of a protein in samples corresponding to the “C−D” condition. All the biological replicate (N=3) are presented in the heatmaps explicitly.

## Notes

### Competing Interest Statement

The authors have declared no competing interest.

## References

(1) Winterbourn, C. Biological Chemistry of Reactive Oxygen Species. In Encyclopedia of Radicals in Chemistry Biology & Materials; 2012; Vol. 3. 10.1002/9781119953678.rad077.

(2) Sies, H. Hydrogen Peroxide as a Central Redox Signaling Molecule in Physiological Oxidative Stress: Oxidative Eustress. Redox Biol. 2017, 11, 613– 619. 10.1016/j.redox.2016.12.035.

(3) Brieger, K.; Schiavone, S.; Miller, F. J.; Krause, K.-H. Reactive Oxygen Species: From Health to Disease. Swiss Med. Wkly. 2012, 142, w13659. 10.4414/smw.2012.13659.

(4) Niethammer, P.; Grabher, C.; Look, A. T.; Mitchison, T. J. A Tissue-Scale Gradient of Hydrogen Peroxide Mediates Rapid Wound Detection in Zebrafish. Nature 2009, 459 (7249), 996–999. 10.1038/nature08119.

(5) Woolley, J. F.; Corcoran, A.; Groeger, G.; Landry, W. D.; Cotter, T. G. Redox-Regulated Growth Factor Survival Signaling. Antioxid. Redox Signal. 2013, 19 (15), 1815–1827. 10.1089/ars.2012.5028.

(6) Winterbourn, C. C. The Biological Chemistry of Hydrogen Peroxide. Methods Enzymol. 2013, 528, 3–25. 10.1016/B978-0-12-405881-1.00001-X.

(7) Mishina, N. M.; Tyurin-Kuzmin, P. A.; Markvicheva, K. N.; Vorotnikov, A. V.; Tkachuk, V. A.; Laketa, V.; Schultz, C.; Lukyanov, S.; Belousov, V. V. Does Cellular Hydrogen Peroxide Diffuse or Act Locally? Antioxid. Redox Signal. 2011, 14 (1), 1–7. 10.1089/ars.2010.3539.

(8) Bestetti, S.; Galli, M.; Sorrentino, I.; Pinton, P.; Rimessi, A.; Sitia, R.; Medraño-Fernandez, I. Human Aquaporin-11 Guarantees Efficient Transport of H2O2 across the Endoplasmic Reticulum Membrane. Redox Biol. 2020, 28, 101326. 10.1016/j.redox.2019.101326.

(9) Joseph, S. K.; Booth, D. M.; Young, M. P.; Hajnóczky, G. Redox Regulation of ER and Mitochondrial Ca2+ Signaling in Cell Survival and Death. Cell Calcium 2019, 79, 89–97. 10.1016/j.ceca.2019.02.006.

(10) Lukosz, M.; Jakob, S.; Büchner, N.; Zschauer, T.-C.; Altschmied, J.; Haendeler, J. Nuclear Redox Signaling. Antioxid. Redox Signal. 2010, 12 (6), 713–742. 10.1089/ars.2009.2609.

(11) Malinouski, M.; Zhou, Y.; Belousov, V. V.; Hatfield, D. L.; Gladyshev, V. N. Hydrogen Peroxide Probes Directed to Different Cellular Compartments. PloS One 2011, 6 (1), e14564. 10.1371/journal.pone.0014564.

(12) Roscoe, J. M.; Sevier, C. S. Pathways for Sensing and Responding to Hydrogen Peroxide at the Endoplasmic Reticulum. Cells 2020, 9 (10), 2314. 10.3390/cells9102314.

(13) Sun, W.; Dai, L.; Yu, H.; Puspita, B.; Zhao, T.; Li, F.; Tan, J. L.; Lim, Y. T.; Chen, M. W.; Sobota, R. M.; Tenen, D. G.; Prabhu, N.; Nordlund, P. Monitoring Structural Modulation of Redox-Sensitive Proteins in Cells with MS-CETSA. Redox Biol. 2019, 24, 101168. 10.1016/j.redox.2019.101168.

(14) Fransen, M.; Lismont, C. Redox Signaling from and to Peroxisomes: Progress, Challenges, and Prospects. Antioxid. Redox Signal. 2019, 30 (1), 95–112. 10.1089/ars.2018.7515.

(15) Bulina, M. E.; Chudakov, D. M.; Britanova, O. V.; Yanushevich, Y. G.; Staroverov, D. B.; Chepurnykh, T. V.; Merzlyak, E. M.; Shkrob, M. A.; Lukyanov, S.; Lukyanov, K. A. A Genetically Encoded Photosensitizer. Nat. Biotechnol. 2006, 24 (1), 95–99. 10.1038/nbt1175.

(16) Sarkisyan, K. S.; Zlobovskaya, O. A.; Gorbachev, D. A.; Bozhanova, N. G.; Sharonov, G. V.; Staroverov, D. B.; Egorov, E. S.; Ryabova, A. V.; Solntsev, K. M.; Mishin, A. S.; Lukyanov, K. A. KillerOrange, a Genetically Encoded Photosensitizer Activated by Blue and Green Light. PloS One 2015, 10 (12), e0145287. 10.1371/journal.pone.0145287.

(17) Sies, H.; Belousov, V. V.; Chandel, N. S.; Davies, M. J.; Jones, D. P.; Mann, G. E.; Murphy, M. P.; Yamamoto, M.; Winterbourn, C. Defining Roles of Specific Reactive Oxygen Species (ROS) in Cell Biology and Physiology. Nat. Rev. Mol. Cell Biol. 2022, 23 (7), 499–515. 10.1038/s41580-022-00456-z.

(18) Pollegioni, L.; Piubelli, L.; Sacchi, S.; Pilone, M. S.; Molla, G. Physiological Functions of D-Amino Acid Oxidases: From Yeast to Humans. Cell. Mol. Life Sci. CMLS 2007, 64 (11), 1373–1394. 10.1007/s00018-007-6558-4.

(19) Day, N. J.; Gaffrey, M. J.; Qian, W.-J. Stoichiometric Thiol Redox Proteomics for Quantifying Cellular Responses to Perturbations. Antioxid. Basel Switz. 2021, 10 (3), 499. 10.3390/antiox10030499.

(20) Shi, Y.; Carroll, K. S. Activity-Based Sensing for Site-Specific Proteomic Analysis of Cysteine Oxidation. Acc. Chem. Res. 2020, 53 (1), 20–31. 10.1021/acs.accounts.9b00562.

(21) Fu, L.; Li, Z.; Liu, K.; Tian, C.; He, J.; He, J.; He, F.; Xu, P.; Yang, J. A Quantitative Thiol Reactivity Profiling Platform to Analyze Redox and Electrophile Reactive Cysteine Proteomes. Nat. Protoc. 2020, 15 (9), 2891–2919. 10.1038/s41596-020-0352-2.

(22) Guo, J.; Nguyen, A. Y.; Dai, Z.; Su, D.; Gaffrey, M. J.; Moore, R. J.; Jacobs, J. M.; Monroe, M. E.; Smith, R. D.; Koppenaal, D. W.; Pakrasi, H. B.; Qian, W.-J. Proteome-Wide Light/Dark Modulation of Thiol Oxidation in Cyanobacteria Revealed by Quantitative Site-Specific Redox Proteomics. Mol. Cell. Proteomics MCP 2014, 13 (12), 3270–3285. 10.1074/mcp.M114.041160.

(23) van der Reest, J.; Lilla, S.; Zheng, L.; Zanivan, S.; Gottlieb, E. Proteome-Wide Analysis of Cysteine Oxidation Reveals Metabolic Sensitivity to Redox Stress. Nat. Commun. 2018, 9 (1), 1581. 10.1038/s41467-018-04003-3.

(24) Xiao, H.; Jedrychowski, M. P.; Schweppe, D. K.; Huttlin, E. L.; Yu, Q.; Heppner, D. E.; Li, J.; Long, J.; Mills, E. L.; Szpyt, J.; He, Z.; Du, G.; Garrity, R.; Reddy, A.; Vaites, L. P.; Paulo, J. A.; Zhang, T.; Gray, N. S.; Gygi, S. P.; Chouchani, E. T. A Quantitative Tissue-Specific Landscape of Protein Redox Regulation during Aging. Cell 2020, 180 (5), 968–983.e24. 10.1016/j.cell.2020.02.012.

(25) Mermelekas, G.; Makridakis, M.; Koeck, T.; Vlahou, A. Redox Proteomics: From Residue Modifications to Putative Biomarker Identification by Gel– and LC-MS-Based Approaches. Expert Rev. Proteomics 2013, 10 (6), 537–549. 10.1586/14789450.2013.855611.

(26) Martinez Molina, D.; Jafari, R.; Ignatushchenko, M.; Seki, T.; Larsson, E. A.; Dan, C.; Sreekumar, L.; Cao, Y.; Nordlund, P. Monitoring Drug Target Engagement in Cells and Tissues Using the Cellular Thermal Shift Assay. Science 2013, 341 (6141), 84–87. 10.1126/science.1233606.

(27) Savitski, M. M.; Reinhard, F. B. M.; Franken, H.; Werner, T.; Savitski, M. F.; Eberhard, D.; Martinez Molina, D.; Jafari, R.; Dovega, R. B.; Klaeger, S.; Kuster, B.; Nordlund, P.; Bantscheff, M.; Drewes, G. Tracking Cancer Drugs in Living Cells by Thermal Profiling of the Proteome. Science 2014, 346 (6205), 1255784. 10.1126/science.1255784.

(28) Mateus, A.; Kurzawa, N.; Becher, I.; Sridharan, S.; Helm, D.; Stein, F.; Typas, A.; Savitski, M. M. Thermal Proteome Profiling for Interrogating Protein Interactions. Mol. Syst. Biol. 2020, 16 (3), e9232. 10.15252/msb.20199232.

(29) Matlashov, M. E.; Belousov, V. V.; Enikolopov, G. How Much H(2)O(2) Is Produced by Recombinant D-Amino Acid Oxidase in Mammalian Cells? Antioxid. Redox Signal. 2014, 20 (7), 1039–1044. 10.1089/ars.2013.5618.

(30) Mishina, N. M.; Bogdanova, Y. A.; Ermakova, Y. G.; Panova, A. S.; Kotova, D. A.; Bilan, D. S.; Steinhorn, B.; Arnér, E. S. J.; Michel, T.; Belousov, V. V. Which Antioxidant System Shapes Intracellular H2O2 Gradients? Antioxid. Redox Signal. 2019, 31 (9), 664–670. 10.1089/ars.2018.7697.

(31) Mehmeti, I.; Lortz, S.; Lenzen, S. The H2O2-Sensitive HyPer Protein Targeted to the Endoplasmic Reticulum as a Mirror of the Oxidizing Thiol-Disulfide Milieu. Free Radic. Biol. Med. 2012, 53 (7), 1451–1458. 10.1016/j.freeradbiomed.2012.08.010.

(32) Saei, A. A.; Gullberg, H.; Sabatier, P.; Beusch, C. M.; Johansson, K.; Lundgren, B.; Arvidsson, P. I.; Arnér, E. S. J.; Zubarev, R. A. Comprehensive Chemical Proteomics for Target Deconvolution of the Redox Active Drug Auranofin. Redox Biol. 2020, 32, 101491. 10.1016/j.redox.2020.101491.

(33) Tan, C. S. H.; Go, K. D.; Bisteau, X.; Dai, L.; Yong, C. H.; Prabhu, N.; Ozturk, M. B.; Lim, Y. T.; Sreekumar, L.; Lengqvist, J.; Tergaonkar, V.; Kaldis, P.; Sobota, R. M.; Nordlund, P. Thermal Proximity Coaggregation for System-Wide Profiling of Protein Complex Dynamics in Cells. Science 2018, 359 (6380), 1170–1177. 10.1126/science.aan0346.

(34) Becher, I.; Andrés-Pons, A.; Romanov, N.; Stein, F.; Schramm, M.; Baudin, F.; Helm, D.; Kurzawa, N.; Mateus, A.; Mackmull, M.-T.; Typas, A.; Müller, C. W.; Bork, P.; Beck, M.; Savitski, M. M. Pervasive Protein Thermal Stability Variation during the Cell Cycle. Cell 2018, 173 (6), 1495–1507.e18. 10.1016/j.cell.2018.03.053.

(35) Becher, I.; Werner, T.; Doce, C.; Zaal, E. A.; Tögel, I.; Khan, C. A.; Rueger, A.; Muelbaier, M.; Salzer, E.; Berkers, C. R.; Fitzpatrick, P. F.; Bantscheff, M.; Savitski, M. M. Thermal Profiling Reveals Phenylalanine Hydroxylase as an Off-Target of Panobinostat. Nat. Chem. Biol. 2016, 12 (11), 908–910. 10.1038/nchembio.2185.

(36) Clemens, M. J. Initiation Factor eIF2 Alpha Phosphorylation in Stress Responses and Apoptosis. Prog. Mol. Subcell. Biol. 2001, 27, 57–89. 10.1007/978-3-662-09889-9_3.

(37) Grant, C. M. Regulation of Translation by Hydrogen Peroxide. Antioxid. Redox Signal. 2011, 15 (1), 191–203. 10.1089/ars.2010.3699.

(38) Shenton, D.; Smirnova, J. B.; Selley, J. N.; Carroll, K.; Hubbard, S. J.; Pavitt, G. D.; Ashe, M. P.; Grant, C. M. Global Translational Responses to Oxidative Stress Impact upon Multiple Levels of Protein Synthesis. J. Biol. Chem. 2006, 281 (39), 29011–29021. 10.1074/jbc.M601545200.

(39) Halbach, F.; Reichelt, P.; Rode, M.; Conti, E. The Yeast Ski Complex: Crystal Structure and RNA Channeling to the Exosome Complex. Cell 2013, 154 (4), 814–826. 10.1016/j.cell.2013.07.017.

(40) Zinder, J. C.; Lima, C. D. Targeting RNA for Processing or Destruction by the Eukaryotic RNA Exosome and Its Cofactors. Genes Dev. 2017, 31 (2), 88–100. 10.1101/gad.294769.116.

(41) van Hoof, A.; Frischmeyer, P. A.; Dietz, H. C.; Parker, R. Exosome-Mediated Recognition and Degradation of mRNAs Lacking a Termination Codon. Science 2002, 295 (5563), 2262–2264. 10.1126/science.1067272.

(42) Jamar, N. H.; Kritsiligkou, P.; Grant, C. M. The Non-Stop Decay mRNA Surveillance Pathway Is Required for Oxidative Stress Tolerance. Nucleic Acids Res. 2017, 45 (11), 6881–6893. 10.1093/nar/gkx306.

(43) Saito, S.; Hosoda, N.; Hoshino, S. The Hbs1-Dom34 Protein Complex Functions in Non-Stop mRNA Decay in Mammalian Cells. J. Biol. Chem. 2013, 288 (24), 17832–17843. 10.1074/jbc.M112.448977.

(44) Heath, C. G.; Viphakone, N.; Wilson, S. A. The Role of TREX in Gene Expression and Disease. Biochem. J. 2016, 473 (19), 2911–2935. 10.1042/BCJ20160010.

(45) Jimeno, S.; Aguilera, A. The THO Complex as a Key mRNP Biogenesis Factor in Development and Cell Differentiation. J. Biol. 2010, 9 (1), 6. 10.1186/jbiol217.

(46) Morris, K. J.; Corbett, A. H. The Polyadenosine RNA-Binding Protein ZC3H14 Interacts with the THO Complex and Coordinately Regulates the Processing of Neuronal Transcripts. Nucleic Acids Res. 2018, 46 (13), 6561–6575. 10.1093/nar/gky446.

(47) Kelly, S. M.; Leung, S. W.; Pak, C.; Banerjee, A.; Moberg, K. H.; Corbett, A. H. A Conserved Role for the Zinc Finger Polyadenosine RNA Binding Protein, ZC3H14, in Control of Poly(A) Tail Length. RNA N. Y. N 2014, 20 (5), 681–688. 10.1261/rna.043984.113.

(48) Soucek, S.; Zeng, Y.; Bellur, D. L.; Bergkessel, M.; Morris, K. J.; Deng, Q.; Duong, D.; Seyfried, N. T.; Guthrie, C.; Staley, J. P.; Fasken, M. B.; Corbett, A. H. The Evolutionarily-Conserved Polyadenosine RNA Binding Protein, Nab2, Cooperates with Splicing Machinery to Regulate the Fate of Pre-mRNA. Mol. Cell. Biol. 2016, 36 (21), 2697–2714. 10.1128/MCB.00402-16.

(49) Marfatia, K. A.; Crafton, E. B.; Green, D. M.; Corbett, A. H. Domain Analysis of the Saccharomyces Cerevisiae Heterogeneous Nuclear Ribonucleoprotein, Nab2p. Dissecting the Requirements for Nab2p-Facilitated Poly(A) RNA Export. J. Biol. Chem. 2003, 278 (9), 6731–6740. 10.1074/jbc.M207571200.

(50) Alqawlaq, S.; Livne-Bar, I.; Williams, D.; D’Ercole, J.; Leung, S. W.; Chan, D.; Tuccitto, A.; Datti, A.; Wrana, J. L.; Corbett, A. H.; Schmitt-Ulms, G.; Sivak, J. M. An Endogenous PI3K Interactome Promoting Astrocyte-Mediated Neuroprotection Identifies a Novel Association with RNA-Binding Protein ZC3H14. J. Biol. Chem. 2021, 296, 100118. 10.1074/jbc.RA120.015389.

(51) Griaud, F.; Pierce, A.; Gonzalez Sanchez, M. B.; Scott, M.; Abraham, S. A.; Holyoake, T. L.; Tran, D. D. H.; Tamura, T.; Whetton, A. D. A Pathway from Leukemogenic Oncogenes and Stem Cell Chemokines to RNA Processing via THOC5. Leukemia 2013, 27 (4), 932–940. 10.1038/leu.2012.283.

(52) Sims, R. J.; Mandal, S. S.; Reinberg, D. Recent Highlights of RNA-Polymerase-II-Mediated Transcription. Curr. Opin. Cell Biol. 2004, 16 (3), 263–271. 10.1016/j.ceb.2004.04.004.

(53) Heine, G. F.; Horwitz, A. A.; Parvin, J. D. Multiple Mechanisms Contribute to Inhibit Transcription in Response to DNA Damage. J. Biol. Chem. 2008, 283 (15), 9555–9561. 10.1074/jbc.M707700200.

(54) Nilson, K. A.; Lawson, C. K.; Mullen, N. J.; Ball, C. B.; Spector, B. M.; Meier, J. L.; Price, D. H. Oxidative Stress Rapidly Stabilizes Promoter-Proximal Paused Pol II across the Human Genome. Nucleic Acids Res. 2017, 45 (19), 11088–11105. 10.1093/nar/gkx724.

(55) Quirós, P. M.; Langer, T.; López-Otín, C. New Roles for Mitochondrial Proteases in Health, Ageing and Disease. Nat. Rev. Mol. Cell Biol. 2015, 16 (6), 345–359. 10.1038/nrm3984.

(56) Parker, M. W.; Botchan, M. R.; Berger, J. M. Mechanisms and Regulation of DNA Replication Initiation in Eukaryotes. Crit. Rev. Biochem. Mol. Biol. 2017, 52 (2), 107–144. 10.1080/10409238.2016.1274717.

(57) Mulvaney, K. M.; Matson, J. P.; Siesser, P. F.; Tamir, T. Y.; Goldfarb, D.; Jacobs, T. M.; Cloer, E. W.; Harrison, J. S.; Vaziri, C.; Cook, J. G.; Major, M. B. Identification and Characterization of MCM3 as a Kelch-like ECH-Associated Protein 1 (KEAP1) Substrate. J. Biol. Chem. 2016, 291 (45), 23719–23733. 10.1074/jbc.M116.729418.

(58) Tamberg, N.; Tahk, S.; Koit, S.; Kristjuhan, K.; Kasvandik, S.; Kristjuhan, A.; Ilves, I. Keap1-MCM3 Interaction Is a Potential Coordinator of Molecular Machineries of Antioxidant Response and Genomic DNA Replication in Metazoa. Sci. Rep. 2018, 8 (1), 12136. 10.1038/s41598-018-30562-y.

(59) Li, Z.; Xu, X. Post-Translational Modifications of the Mini-Chromosome Maintenance Proteins in DNA Replication. Genes 2019, 10 (5), 331. 10.3390/genes10050331.

(60) Raman, D.; Pervaiz, S. Redox Inhibition of Protein Phosphatase PP2A: Potential Implications in Oncogenesis and Its Progression. Redox Biol. 2019, 27, 101105. 10.1016/j.redox.2019.101105.

(61) Salmeen, A.; Andersen, J. N.; Myers, M. P.; Meng, T.-C.; Hinks, J. A.; Tonks, N. K.; Barford, D. Redox Regulation of Protein Tyrosine Phosphatase 1B Involves a Sulphenyl-Amide Intermediate. Nature 2003, 423 (6941), 769–773. 10.1038/nature01680.

(62) Schwertassek, U.; Haque, A.; Krishnan, N.; Greiner, R.; Weingarten, L.; Dick, T. P.; Tonks, N. K. Reactivation of Oxidized PTP1B and PTEN by Thioredoxin 1. FEBS J. 2014, 281 (16), 3545–3558. 10.1111/febs.12898.

(63) Ashburner, M.; Ball, C. A.; Blake, J. A.; Botstein, D.; Butler, H.; Cherry, J. M.; Davis, A. P.; Dolinski, K.; Dwight, S. S.; Eppig, J. T.; Harris, M. A.; Hill, D. P.; Issel-Tarver, L.; Kasarskis, A.; Lewis, S.; Matese, J. C.; Richardson, J. E.; Ringwald, M.; Rubin, G. M.; Sherlock, G. Gene Ontology: Tool for the Unification of Biology. The Gene Ontology Consortium. Nat. Genet. 2000, 25 (1), 25–29. 10.1038/75556.

(64) Gene Ontology Consortium. The Gene Ontology Resource: Enriching a GOld Mine. Nucleic Acids Res. 2021, 49 (D1), D325–D334. 10.1093/nar/gkaa1113.

(65) Richarme, G.; Mihoub, M.; Dairou, J.; Bui, L. C.; Leger, T.; Lamouri, A. Parkinsonism-Associated Protein DJ-1/Park7 Is a Major Protein Deglycase That Repairs Methylglyoxal– and Glyoxal-Glycated Cysteine, Arginine, and Lysine Residues. J. Biol. Chem. 2015, 290 (3), 1885–1897. 10.1074/jbc.M114.597815.

(66) Chen, J.; Li, L.; Chin, L.-S. Parkinson Disease Protein DJ-1 Converts from a Zymogen to a Protease by Carboxyl-Terminal Cleavage. Hum. Mol. Genet. 2010, 19 (12), 2395–2408. 10.1093/hmg/ddq113.

(67) Mencke, P.; Boussaad, I.; Romano, C. D.; Kitami, T.; Linster, C. L.; Krüger, R. The Role of DJ-1 in Cellular Metabolism and Pathophysiological Implications for Parkinson’s Disease. Cells 2021, 10 (2), 347. 10.3390/cells10020347.

(68) Matassa, D. S.; Arzeni, D.; Landriscina, M.; Esposito, F. ER Stress Protection in Cancer Cells: The Multifaceted Role of the Heat Shock Protein TRAP1. Cell Pathol. 2014, 1 (1), 40–48. 10.2478/ersc-2014-0003.

(69) Masgras, I.; Sanchez-Martin, C.; Colombo, G.; Rasola, A. The Chaperone TRAP1 As a Modulator of the Mitochondrial Adaptations in Cancer Cells. Front. Oncol. 2017, 7, 58. 10.3389/fonc.2017.00058.

(70) Stankovic-Valentin, N.; Melchior, F. Control of SUMO and Ubiquitin by ROS: Signaling and Disease Implications. Mol. Aspects Med. 2018, 63, 3–17. 10.1016/j.mam.2018.07.002.

(71) Braun, H. P.; Schmitz, U. K. The Mitochondrial Processing Peptidase. Int. J. Biochem. Cell Biol. 1997, 29 (8–9), 1043–1045. 10.1016/s1357-2725(97)00032-0.

(72) Malty, R. H.; Aoki, H.; Kumar, A.; Phanse, S.; Amin, S.; Zhang, Q.; … & Babu, M. A Map of Human Mitochondrial Protein Interactions Linked to Neurodegeneration Reveals New Mechanisms of Redox Homeostasis and NF-κB Signaling. Cell Systems, 2017, 5 (6), 564 – 577.e12. 10.1016/j.cels.2017.10.010.

(73) Szklarczyk, D.; Kirsch, R.; Koutrouli, M.; Nastou, K.; Mehryary, F.; Hachilif, R.; … & Von Mering, C. The STRING database in 2023: protein–protein association networks and functional enrichment analyses for any sequenced genome of interest. Nucleic Acids Res. 2023, 51(D1): D638–646. 10.1093/nar/gkac1000.

(74) Li, H. M.; Niki, T.; Taira, T.; Iguchi-Ariga, S. M. M.; Ariga, H. Association of DJ-1 with Chaperones and Enhanced Association and Colocalization with Mitochondrial Hsp70 by Oxidative Stress. Free Radic. Res. 2005, 39 (10), 1091–1099. 10.1080/10715760500260348.

(75) Zhou, T.-T.; Wang, X.-Y.; Huang, J.; Deng, Y.-Z.; Qiu, L.-J.; Liu, H.-Y.; Xu, X.-W.; Ma, Z.-X.; Tang, L.; Chen, H.-P. Mitochondrial Translocation of DJ-1 Is Mediated by Grp75: Implication in Cardioprotection of Resveratrol Against Hypoxia/Reoxygenation-Induced Oxidative Stress. J. Cardiovasc. Pharmacol. 2020, 75 (4), 305–313. 10.1097/FJC.0000000000000805.

(76) Canet-Avilés, R. M.; Wilson, M. A.; Miller, D. W.; Ahmad, R.; McLendon, C.; Bandyopadhyay, S.; Baptista, M. J.; Ringe, D.; Petsko, G. A.; Cookson, M. R. The Parkinson’s Disease Protein DJ-1 Is Neuroprotective Due to Cysteine-Sulfinic Acid-Driven Mitochondrial Localization. Proc. Natl. Acad. Sci. U. S. A. 2004, 101 (24), 9103–9108. 10.1073/pnas.0402959101.

(77) Basso, V.; Marchesan, E.; Ziviani, E. A Trio Has Turned into a Quartet: DJ-1 Interacts with the IP3R-Grp75-VDAC Complex to Control ER-Mitochondria Interaction. Cell Calcium 2020, 87, 102186. 10.1016/j.ceca.2020.102186.

(78) Görlach, A.; Bertram, K.; Hudecova, S.; Krizanova, O. Calcium and ROS: A Mutual Interplay. Redox Biol. 2015, 6, 260–271. 10.1016/j.redox.2015.08.010.

(79) Lee, D.-H.; Kim, D.; Kim, S. T.; Jeong, S.; Kim, J. L.; Shim, S. M.; Heo, A. J.; Song, X.; Guo, Z. S.; Bartlett, D. L.; Oh, S. C.; Lee, J.; Saito, Y.; Kim, B. Y.; Kwon, Y. T.; Lee, Y. J. PARK7 Modulates Autophagic Proteolysis through Binding to the N-Terminally Arginylated Form of the Molecular Chaperone HSPA5. Autophagy 2018, 14 (11), 1870–1885. 10.1080/15548627.2018.1491212.

(80) Wilson, M. A. The Role of Cysteine Oxidation in DJ-1 Function and Dysfunction. Antioxid. Redox Signal. 2011, 15 (1), 111–122. 10.1089/ars.2010.3481.

(81) Roberts, P. J.; Der, C. J. Targeting the Raf-MEK-ERK Mitogen-Activated Protein Kinase Cascade for the Treatment of Cancer. Oncogene 2007, 26 (22), 3291–3310. 10.1038/sj.onc.1210422.

(82) Caunt, C. J.; Sale, M. J.; Smith, P. D.; Cook, S. J. MEK1 and MEK2 Inhibitors and Cancer Therapy: The Long and Winding Road. Nat. Rev. Cancer 2015, 15 (10), 577–592. 10.1038/nrc4000.

(83) Obsilova, V.; Obsil, T. The 14-3-3 Proteins as Important Allosteric Regulators of Protein Kinases. Int. J. Mol. Sci. 2020, 21 (22), 8824. 10.3390/ijms21228824.

(84) Bossis, G.; Melchior, F. Regulation of SUMOylation by Reversible Oxidation of SUMO Conjugating Enzymes. Mol. Cell 2006, 21 (3), 349–357. 10.1016/j.molcel.2005.12.019.

(85) Saitoh, H.; Hinchey, J. Functional Heterogeneity of Small Ubiquitin-Related Protein Modifiers SUMO-1 versus SUMO-2/3. J. Biol. Chem. 2000, 275 (9), 6252–6258. 10.1074/jbc.275.9.6252.

(86) Lu, J.; Holmgren, A. The Thioredoxin Antioxidant System. Free Radic. Biol. Med. 2014, 66, 75–87. 10.1016/j.freeradbiomed.2013.07.036.

(87) Rhee, S. G.; Woo, H. A.; Kil, I. S.; Bae, S. H. Peroxiredoxin Functions as a Peroxidase and a Regulator and Sensor of Local Peroxides. J. Biol. Chem. 2012, 287 (7), 4403–4410. 10.1074/jbc.R111.283432.

(88) Heo, S.; Kim, S.; Kang, D. The Role of Hydrogen Peroxide and Peroxiredoxins throughout the Cell Cycle. Antioxid. Basel Switz. 2020, 9 (4), 280. 10.3390/antiox9040280.

(89) Topf, U.; Suppanz, I.; Samluk, L.; Wrobel, L.; Böser, A.; Sakowska, P.; Knapp, B.; Pietrzyk, M. K.; Chacinska, A.; Warscheid, B. Quantitative Proteomics Identifies Redox Switches for Global Translation Modulation by Mitochondrially Produced Reactive Oxygen Species. Nat. Commun. 2018, 9 (1), 324. 10.1038/s41467-017-02694-8.

(90) MacKenzie, J. A.; Payne, R. M. Mitochondrial Protein Import and Human Health and Disease. Biochim. Biophys. Acta 2007, 1772 (5), 509–523. 10.1016/j.bbadis.2006.12.002.

(91) Beyer, H. M.; Gonschorek, P.; Samodelov, S. L.; Meier, M.; Weber, W.; Zurbriggen, M. D. AQUA Cloning: A Versatile and Simple Enzyme-Free Cloning Approach. PloS One 2015, 10 (9), e0137652. 10.1371/journal.pone.0137652.

(92) Hughes, J. R.; Roberts, N.; McGowan, S.; Hay, D.; Giannoulatou, E.; Lynch, M.; De Gobbi, M.; Taylor, S.; Gibbons, R.; Higgs, D. R. Analysis of Hundreds of Cis-Regulatory Landscapes at High Resolution in a Single, High-Throughput Experiment. Nat. Genet. 2014, 46 (2), 205–212. 10.1038/ng.2871.

(93) Mateus, A.; Hevler, J.; Bobonis, J.; Kurzawa, N.; Shah, M.; Mitosch, K.; Goemans, C. V.; Helm, D.; Stein, F.; Typas, A.; Savitski, M. M. The Functional Proteome Landscape of Escherichia Coli. Nature 2020, 588 (7838), 473–478. 10.1038/s41586-020-3002-5.

(94) Franken, H.; Mathieson, T.; Childs, D.; Sweetman, G. M. A.; Werner, T.; Tögel, I.; Doce, C.; Gade, S.; Bantscheff, M.; Drewes, G.; Reinhard, F. B. M.; Huber, W.; Savitski, M. M. Thermal Proteome Profiling for Unbiased Identification of Direct and Indirect Drug Targets Using Multiplexed Quantitative Mass Spectrometry. Nat. Protoc. 2015, 10 (10), 1567–1593. 10.1038/nprot.2015.101.

(95) Huber, W.; von Heydebreck, A.; Sültmann, H.; Poustka, A.; Vingron, M. Variance Stabilization Applied to Microarray Data Calibration and to the Quantification of Differential Expression. Bioinforma. Oxf. Engl. 2002, 18 *Suppl 1*, S96–104. 10.1093/bioinformatics/18.suppl_1.s96.

(96) Strimmer, K. Fdrtool: A Versatile R Package for Estimating Local and Tail Area-Based False Discovery Rates. Bioinforma. Oxf. Engl. 2008, 24 (12), 1461–1462. 10.1093/bioinformatics/btn209.

(97) Klopfenstein, D. V.; Zhang, L.; Pedersen, B. S.; Ramírez, F.; Warwick Vesztrocy, A.; Naldi, A.; Mungall, C. J.; Yunes, J. M.; Botvinnik, O.; Weigel, M.; Dampier, W.; Dessimoz, C.; Flick, P.; Tang, H. GOATOOLS: A Python Library for Gene Ontology Analyses. Sci. Rep. 2018, 8 (1), 10872. 10.1038/s41598-018-28948-z.

(98) Luo, T.; Pueyo, J. M.; Wahni, K.; Yvanoff, C.; Lazar, T.; Pyr Dit Ruys, S.; Vertommen, D.; Ezeriņa, D.; Messens, J. Thiol-Disulphide Independent in-Cell Trapping for the Identification of Peroxiredoxin 2 Interactors. Redox Biol. 2021, 46, 102066. 10.1016/j.redox.2021.102066.

